# A key role for Canoe’s intrinsically disordered region in linking cell junctions to the cytoskeleton

**DOI:** 10.1101/2025.05.10.653259

**Authors:** Corbin C. Jensen, Noah J. Gurley, Avery J. Mathias, Leah R. Wolfsberg, Yufei Xiao, Zixi Zhou, Kevin C. Slep, Mark Peifer

## Abstract

Adherens junctions are key to tissue architecture, mediating robust yet dynamic cell-cell adhesion and, via cytoskeletal linkage, allowing cells to change shape and move. Adherens junctions contain thousands of molecules linked by multivalent interactions of folded protein domains and Intrinsically Disordered Regions (IDRs). One key challenge is defining mechanisms conferring robust linkage and mechanosensing. Drosophila Canoe and mammalian Afadin provide superb entrypoints to explore how their complex protein structures and shared IDRs enable function. We combined genetic, cell biological and biochemical tools to define how Canoe’s IDR functions during morphogenesis. Unlike many of Canoe’s folded domains, the IDR is critical for junctional localization, mechanosensing and function. We took the IDR apart, identifying two conserved stickers in the IDR that directly bind F-actin, separated by less-conserved spacers. Surprisingly, while mutants lacking the IDR die as embryos with morphogenesis defects, no sub-region of the IDR is essential for viability. Instead, IDR stickers and spacers act combinatorially to ensure localization, mechanosensing and function.

## Introduction

Many key cellular functions are carried out by large multi-protein complexes rather than individual proteins. Some form structures known as biomolecular condensates which can contain thousands of molecules linked by multivalent interactions of both folded protein domains and intrinsically disordered regions (IDRs) (Banani et al., 2017). Understanding how these assemble is a key challenge for our field. A premier example is the cell-cell adherens junction (AJ), which joins cells to one another, and, via linkage to the actomyosin cytoskeleton, mediates cell shape change and cell migration. At its core is the cadherin-catenin complex. The extracellular domains of transmembrane cadherins mediate homophilic cell adhesion, while their cytoplasmic tails link to actin via β-catenin (βcat), and α-catenin (αcat). While the connection between AJs and the cytoskeleton was initially thought to rely on this simple linear linkage, we now know it is mediated by a dynamic and multivalent network including many more proteins (Perez-Vale and Peifer, 2020). Further, these proteins provide mechanosensing and mechano-responsiveness, as AJ linkage to actin is strengthened when force is exerted on AJs (Yap et al., 2018).

Textbook diagrams often provide simple pictures of individual cadherins linked to their cytoplasmic partners. However, cadherins assemble in much larger complexes, linking in cis to cadherins in the same cell and in trans to cadherins on neighboring cells. Rather than being arrayed continuously along the apico-lateral membrane, AJ proteins form discrete puncta (Truong Quang et al., 2013). In Drosophila, the Harris lab counted molecules in spot AJs assembling during cellularization (McGill et al., 2009). Bicellular borders have 3-5 puncta, each containing ∼1500 cadherin-catenin complexes and 200 Bazooka (Baz)/Par3 proteins. However, these are not “standardized”; size and protein number per punctum varies and proteins dynamically enter and leave puncta. As cells begin to change shape during gastrulation, puncta increase in number and decrease in apparent brightness (Schmidt et al., 2023). Similar puncta are observed in cultured mammalian cells (e.g. (Choi et al., 2016; Erami et al., 2015; Wu et al., 2015), though their architecture varies widely depending on cell type. This is not unique to AJs—tight junctions and focal contacts also assemble as biomolecular condensates (Citi, 2020; Sun et al., 2022).

A hallmark feature of many cell junction proteins is the presence of long IDRs (Rouaud et al., 2020)). IDRs perform many functions in the assembly of multiprotein complexes. Most are of low sequence complexity, with enrichment of specific subsets of amino acids. They mediate interactions via two broad mechanisms (Choi et al., 2020). First, embedded in IDRs are Short Linear amino acid Motifs (SLiMs), sometimes referred to as “stickers”, that serve as binding sites for other proteins. Some SLiMs fold on their own while others fold upon interaction with a binding partner or bind as extended peptides. These motifs are embedded in unstructured regions that act as “spacers” and can also mediate low affinity interactions. In some cases, these regions confer upon IDRs the ability to phase separate in vitro, and in some models, phase separation also helps drive multiprotein complex assembly in vivo. The IDRs of multiple junctional proteins, including ZO-1 (Beutel et al., 2019), βcat (Zamudio et al., 2019), Par3 (Liu et al., 2020), and Afadin (Kuno et al., 2025) can phase separate. Apart from ZO-1, the roles of this in vivo remains unclear.

Assembling AJs involves multivalent interactions among the many proteins in the AJ network. The Drosophila AJ-cytoskeletal linker Canoe (Cno) and its mammalian ortholog Afadin play key roles in multiple morphogenic movements during embryonic and postembryonic morphogenesis and homeostasis, making them outstanding entry-points for studying AJ assembly and dynamics. Cno regulates diverse processes including apical positioning of AJs, apical constriction of mesodermal cells, cell intercalation and axis elongation, and epithelial sheet migration during dorsal closure and head involution (Boettner et al., 2003; Choi et al., 2013; Sawyer et al., 2011; Sawyer et al., 2009). Cno is mechanosensitive, with recruitment to AJs enhanced under tension (Yu and Zallen, 2020). This stabilizes AJ:cytoskeletal linkage (Perez-Vale et al., 2021). In Cno’s absence, AJ/cytoskeletal connections are disrupted, with AJ gaps and the cytoskeleton broadened under tension. Similar effects are seen in mammalian cells with reduced Afadin (Choi et al., 2016; Sakakibara et al., 2020).

The second reason Cno/Afadin is a superb entry-point is its size and structural complexity. Cno/Afadin is a multidomain protein, with five well-conserved folded protein domains followed by a long IDR (Gurley et al., 2023). This allows for multivalent interactions with other proteins in the AJ network. The two N-terminal Ras-association (RA) domains each bind the membrane-anchored small GTPase Rap1 when it is in its active state, and this in turn “activates” Cno (McParland et al., 2024b). The PDZ domain binds AJ transmembrane proteins including nectins (Takahashi et al., 1999) and E-cadherin (Ecad) (Sawyer et al., 2009). Meanwhile, regions in the IDR bind F-actin (Carminati et al., 2016; Mandai et al., 1997; Sawyer et al., 2009), αcat (Pokutta et al., 2002), and other junctional proteins.

To elucidate the molecular mechanisms underlying Cno function, we are systematically taking it apart, deleting specific domains or regions and assessing their roles. We initially hypothesized Rap1 binding to the RA domains activated Cno and Cno then used its PDZ domain and the so-called F-actin binding (FAB) region of the IDR to link Ecad and actin. We tested this, using CRISPR to generate *cno* mutants cleanly deleting each domain/region (Perez-Vale et al., 2021). Mutants lacking both RA domains die as embryos with severe morphogenesis defects, consistent with our hypothesis. In contrast, neither the PDZ domain nor the FAB region are required for viability to adulthood. Subsequent work revealed that the RA2 and Dilute domains are also individually dispensable (McParland et al., 2024a; McParland et al., 2024b; Perez-Vale et al., 2021).

However, each mutant has reduced function in sensitized assays. This changed our thinking, emphasizing how robustness of AJ-cytoskeletal connections is conferred by a network of proteins linked by multivalent interactions.

While Cno and Afadin share long IDRs, they differ in length (952 vs 802 amino acids), charge, and sequence composition (Gurley et al., 2023). The C-terminal FAB region is modestly conserved (35% sequence identity; (Perez-Vale et al., 2021)), but reciprocal BLAST searches with the proximal IDR revealed “no significant similarity”. Binding sites in Afadin’s IDR for ZO-1 (Ooshio et al., 2010) and Lgn (Carminati et al., 2016) do not appear to be conserved in Cno. We used AlphaFold to identify potential structures in the IDRs (Gurley et al., 2023). Afadin’s IDR has multiple predicted α-helical regions. These include a region known to bind the M-domain of αcat (Maruo et al., 2018; Pokutta et al., 2002), but surprisingly this helix is not conserved in Cno. In contrast, two predicted helices in the middle of both IDRs are conserved (33% identity; (Gurley et al., 2023). This set of two helices in Afadin can bind both the actin-binding domain of αcat and actin filaments, thus stabilizing α-catenin’s interaction with actin (Gong et al., 2025). There also are predicted helices in the FAB region.

The IDRs of both Afadin and Cno bind F-actin (Mandai et al., 1997; Sawyer et al., 2009). Initial studies used fragments covering the C-terminal half of the IDR. Subsequent work narrowed one interacting region in Afadin to the two conserved α-helices in the middle of the IDR along with a more C-terminal helix only found in Afadin (Carminati et al., 2016). The conserved helices alone can co-sediment with actin in the presence of αcat and enhance αcat:actin binding; cryo-EM revealed the structural basis for this (Gong et al., 2025). In contrast, despite the designation of the C-terminal end of the IDR as the “F-actin binding region” (FAB), no one has directly assessed actin binding.

Functional assays of the IDR are limited. Deletion studies suggest a modest role for the FAB in full function of both Afadin (Sakakibara et al., 2018) and Cno (Perez-Vale et al., 2021). Mutations leading to premature stop codons in Cno’s IDR have strong phenotypes, but we could not rule out effects of nonsense-mediated mRNA decay (Gurley et al., 2023). In mammalian cells, Afadin knockdown (Choi et al., 2016) or knockout (Gong et al., 2025; Sakakibara et al., 2020) broaden actin at bicellular and tricellular junctions (TCJs). A mutant protein lacking both the helix mediating binding to the M-domain of αcat and the two conserved helices could not rescue junctional actin architecture, while adding this region alone provided partial rescue (Sakakibara et al., 2020). More limited analysis of an Afadin mutant retaining the FAB but lacking the proximal IDR revealed that it still localizes to AJs but loses apical enrichment and apical junctions are destabilized (Kuno et al., 2025). Here we extended and expanded this analysis by combining genetic, cell biological and biochemical tools to define the function of the IDR and its stickers and spacers in Cno localization and function in vivo during morphogenesis.

## Results

### Cno’s proximal IDR is essential for embryonic viability

We previously assessed the role of the IDR region with the strongest sequence conservation between Drosophila and mammals, referred to as the FAB. The FAB is dispensable for viability and fertility, though sensitized tests revealed it contributes to the robustness of morphogenesis (Perez-Vale et al., 2021). To test the function of the rest of the IDR (the proximal IDR (ProxIDR)), we generated a Cno mutant deleting sequences from the PDZ domain to the beginning of the FAB (Fig 1A; construct details are in Fig S1). This mutant, with an added C-terminal GFP, was engineered into the endogenous *cno* locus, using the platform we developed (Perez-Vale et al., 2021). We refer to this mutant, as *cnoΔProxIDR.* This and all other mutants described below were verified by sequencing genomic DNA, and we used immunoblotting to confirm that appropriately sized and GFP-tagged proteins were produced at roughly wildtype levels (Fig S2A, B).

**Fig 1.**
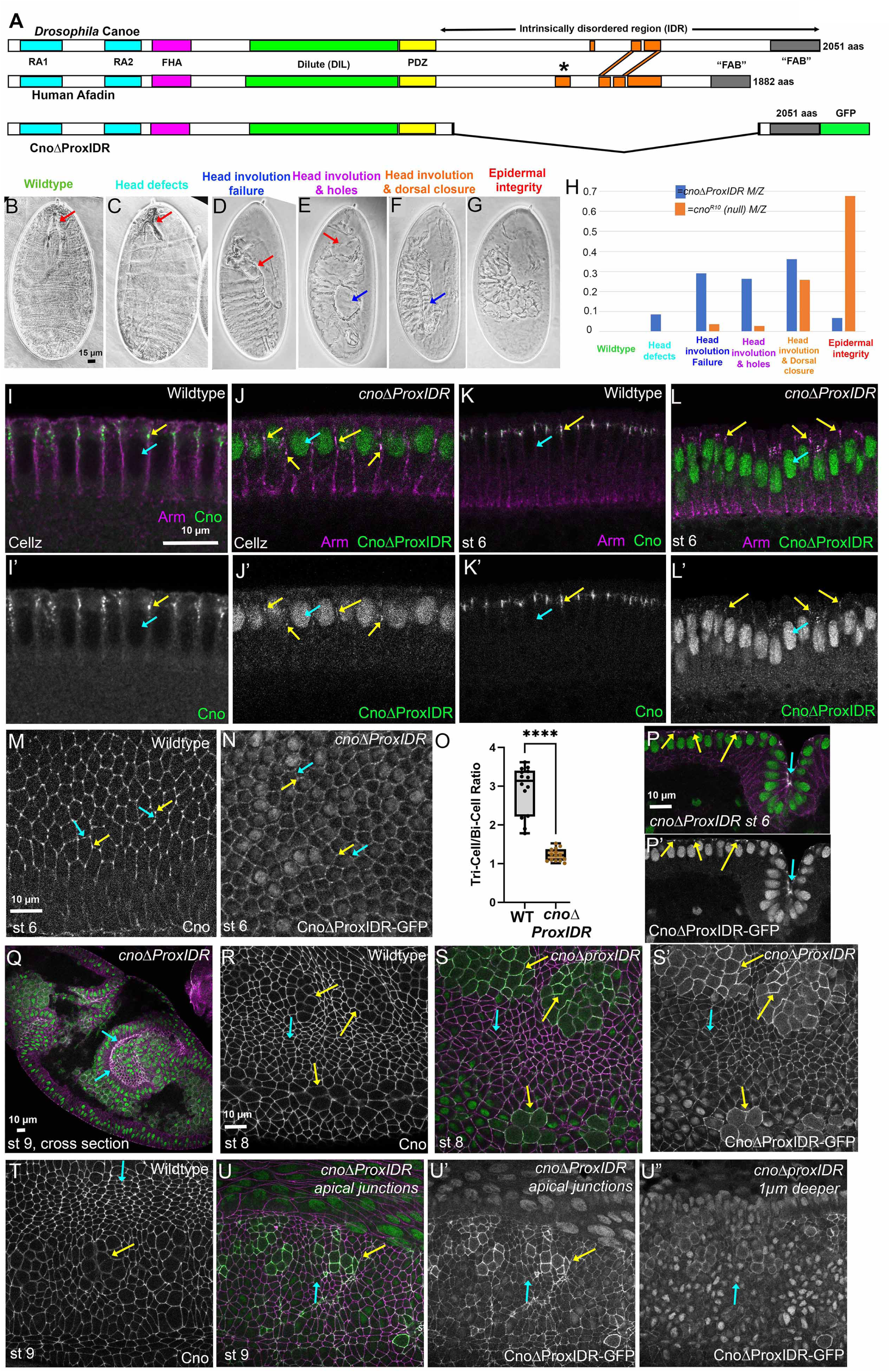
Cno’s IDR is critical for Cno’s role in embryonic morphogenesis and in Cno protein localization. **(A).** Drosophila Cno, Human Afadin, and CnoΔProxIDR. Orange boxes=predicted helices. **(B-G).** Cuticles illustrating different strength phenotypes. B-E. Red arrows indicate head skeleton (B-C) or complete head involution failure (D,E). **(E, F).** Blue arrows=dorsal hole (E) or complete failure of dorsal closure (F). **(G).** Most extreme defects in epidermal integrity. **(H).** Distribution of cuticle phenotypes in different mutants. *cno^R10^/+* data is from (Gurley et al., 2023). **(I-N, P-U).** Embryos, stained to visualize Cno or GFP-tagged CnoΔProxIDR. **(I-L, P).** Cross-sections, apical up, stages indicated. Yellow arrows=junctional localization. Cyan arrows=nuclei. Red arrow=ectodermal fold. **(M, N).** Apical view, level of AJs. Yellow arrows=TCJs, cyan arrows=bicellular junctions**. (O)** Tricellular junction enrichment**. (Q).** Whole embryo. Arrow=enrichment in invaginating midgut. **(S-U).** Stages 8-9. Yellow arrows=cells rounded up for mitosis. Cyan arrows=nondividing cells.

We first examined whether *cnoΔProxIDR* was viable over our null allele, *cno^R2^*. No adult progeny were seen (0/174), thus contrasting with *cnoΔRA2, cnoΔDIL, cnoΔPDZ,* and *cnoΔFAB,* all of which are adult viable (McParland et al., 2024a; McParland et al., 2024b; Perez-Vale et al., 2021). However, unlike null alleles like *cno^R2^*, *cnoΔProxIDR* is not zygotically embryonic lethal (7% lethality; n=755; in the wildtype 3-8% range). To assess the function of CnoΔProxIDR in embryonic morphogenesis, we generated maternal-zygotic mutants, using the FLP/FRT approach (Chou and Perrimon, 1996) to generate females with germlines homozygous for *cnoΔProxIDR* and crossing them to *cnoΔProxIDR/+* males. 50% of the offspring are maternal-zygotic mutants, and we observed 57% embryonic lethality (n=509), consistent with fully penetrant lethality of maternal-zygotic mutants and substantial zygotic rescue. Thus, the IDR is essential for embryonic viability.

### Cno’s proximal IDR plays important roles in embryonic morphogenesis, but some Cno function remains in its absence

As a first assessment of CnoΔProxIDR function in embryonic morphogenesis, we examined larval cuticles of dead embryos. This analysis allows us to assess the completion of multiple morphogenic events requiring Cno function, including dorsal closure, head involution and ventral epidermal integrity (Fig 1B-G).

Maternal/zygotic *cno* null mutants have strong defects in all these events (Fig 1H), with fully penetrant failure of dorsal closure and head involution, and with ∼70% of embryos exhibiting additional defects in epidermal integrity (as in Fig 1G; (Gurley et al., 2023). *cnoΔProxIDR* maternal-zygotic mutants had substantial defects in morphogenesis, but these were not as severe as those of *cno* null mutants. Only 43% of *cnoΔProxIDR* mutants had complete failure of dorsal closure and head involution (phenotypes in Fig 1F+G), while 37% were substantially less severe, with defects in head involution but which completed dorsal closure (as in Fig 1C-E; n=293). Thus, the proximal IDR is important for Cno function in morphogenesis, but some function remains in its absence.

### Deleting Cno’s proximal IDR relocalizes the protein from AJs to the nucleus

In wildtype embryos Cno localizes to nascent AJs as they assemble during cellularization (Fig 1I, yellow arrow; (Bonello et al., 2018; Choi et al., 2013)) and remains localized to apical junctions from the onset of gastrulation throughout the remainder of development (Fig 1K, yellow arrow; (Sawyer et al., 2009). While two of our previous mutant proteins, CnoΔRA and CnoΔRA1, exhibited altered localization during cellularization, all mutant proteins localized to AJs after gastrulation onset. We thus explored localization of CnoΔProxIDR.

The results were quite surprising. During cellularization, CnoΔProxIDR was predominantly localized to nuclei (Fig 1J, cyan arrow). Low levels of CnoΔProxIDR protein accumulated at spot AJs (Fig 1J, yellow arrows), but unlike wildtype Cno (Fig 1I) this junctional localization was not restricted to apical junctional planes. As gastrulation commenced in stage 6, CnoΔProxIDR remained predominantly nuclear, but junctional localization increased (Fig 1L, cyan vs yellow arrows). During germband extension, wildtype Cno localizes to all AJs and is especially enriched at tricellular junctions (TCJs) relative to bicellular junctions (Fig 1M, yellow vs cyan arrows). At this stage, CnoΔProxIDR retains strong nuclear localization along with some junctional enrichment (Fig 1N), but TCJ enrichment relative to levels at bicellular junctions is lost (Fig 1N, yellow vs cyan arrows; quantified in O). Intriguingly, AJ enrichment of CnoΔProxIDR was elevated in cells that were apically constricting to form folds or the posterior midgut invagination (Fig1P, Q, cyan arrows), as we previously observed for wildtype Cno.

During stage 8 cell division commences in the thorax and abdomen. Cells enter mitosis in programmed groups called mitotic domains (Foe, 1989). During mitosis, cells round up and nuclear contents are released into the cytoplasm. The AJ localization of core AJ proteins and Cno is somewhat reduced in cells as they round up relative to cells not undergoing division (Fig 1R, yellow vs cyan arrows), perhaps by simple dilution as junctional perimeter increases. However, we observed a striking change in the localization of CnoΔProxIDR. No longer confined to nuclei, CnoΔProxIDR filled the cytoplasm of dividing cells and became substantially more enriched at the cell cortex relative to neighboring non-mitotic cells (Fig 1S, yellow vs cyan arrows). These features of protein localization continued through the rest of morphogenesis, with weaker AJ localization of CnoΔProxIDR in non-dividing cells and stronger cortical localization in mitotic cells (Fig 1U, cyan vs yellow arrows), in contrast to what we observe with wildtype Cno (Fig 1T). Upon sectioning 1µm deeper, the continuing localization of CnoΔProxIDR to nuclei is apparent (Fig 1U”, arrow). Thus the proximal IDR is critical for effective recruitment/retention of Cno at AJs, as when it is missing the protein substantially re-localizes to nuclei. We explore implications of this in the Discussion.

### *cnoΔProxIDR* mutants have defects in morphogenesis and AJ integrity but these are less severe than those of *cno* null mutants

Next we explored the function of CnoΔProxIDR protein in detail. Cno’s first role in morphogenesis comes at the onset of gastrulation when ventral cells apically constrict and invaginate to become mesoderm (Fig 2A-B). In *cno* null mutants the contractile network of actin and myosin detaches from AJs and invagination stalls, leading to fully penetrant defects in mesoderm invagination and an open ventral furrow (Sawyer et al., 2009). Later some embryos zip up at the midline but defects persist. *cnoΔ*RA mutants also have fully penetrant defects in mesoderm invagination, though defect severity is reduced on average relative to null mutants (Perez-Vale et al., 2021). We examined the success of ventral furrow invagination in *cnoΔProxIDR* mutants, using staining with our C-terminal anti-Cno antibody which does not recognize CnoΔProxIDR, to distinguish maternal zygotic mutants from zygotically-rescued embryos. Many *cnoΔProxIDR* mutants had defects in closing the ventral furrow (Table S1), but these were substantially less penetrant and less severe than seen in *cno* null mutants. 6/31 mutants at stages 6-9 had closed ventral furrows, 10/31 had mild defects in closure (short regions where the furrow was open; Fig 2C), 8/31 had moderate defects (>30% open), and 7/31 had a wide-open ventral furrow (Fig 2D).

**Fig 2.**
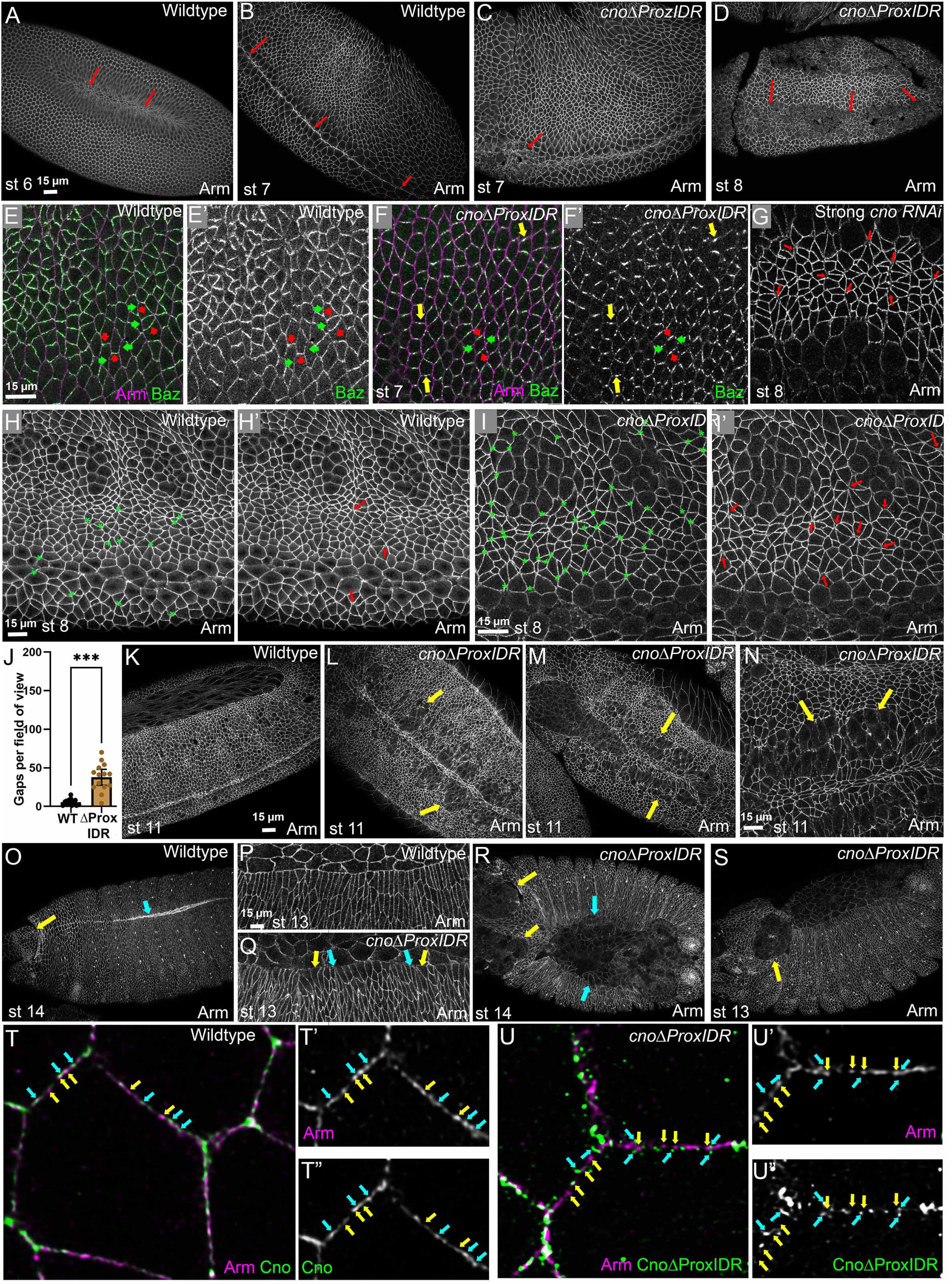
*cnoΔProxIDR* mutants have defects in morphogenesis and junctional integrity, but these are less severe than those of *cno* null mutants. Embryos, anterior left, antigens and stages indicated. **(A-D).** Ventral views, arrows=ventral furrow. **(E-F)** Lateral views. Red arrows=DV borders, green arrows=AP borders, yellow arrows=Baz restricted to center of DV border. **(G-I).** Stage 8. Asterisks=all junctional gaps/disruptions. Arrows=selected gaps. **(J).** Gap quantification. **(K-N).** Stage 11. Arrows=Failure of cells to resume columnar architecture. **(O, P).** Wildtype dorsal closure (cyan arrow) and head involution (yellow arrow). Leading edge cells are relatively uniform in width (P). (Q-S). *cnoΔProxIDR* mutants. **(Q).** Hyper-constricted and splayed open leading edge cells. **(R, S).** Defects in head involution (yellow arrows) and dorsal closure (cyan arrow). **(T, U).** Enhanced resolution imaging of AJs using the Airyscan module. Yellow arrows=junctional puncta enriched for Arm. Cyan arrows=junctional puncta enriched for Cno.

As gastrulation began in *cnoΔProxIDR* mutants, AJs assembled with the βcat homolog Armadillo (Arm) enriched apically but also localized along the lateral interface, thus matching wildtype (Fig 1K vs L). The Par3 ortholog Baz also localizes to apical junctions at this stage. In wildtype, Baz localizes all around the cell but is planar-polarized, with two-fold higher accumulation on dorsal-ventral (DV) borders relative to anterior-posterior (AP) cell borders (Fig 2E, red vs. green arrows; (Zallen and Wieschaus, 2004). However, in both *cno* null and *cnoΔRA* mutants (Perez-Vale et al., 2021; Sawyer et al., 2011), Baz planar polarization is strongly enhanced. This was also true in *cnoΔProxIDR* mutants. Baz localization was relatively enriched on DV cell borders (Fig 2F; red vs. green arrows) and was restricted to the center of the DV borders in some cells (Fig 2F, yellow arrows).

In *cno* null mutants, *cnoΔRA* mutants, or in embryos with strong RNAi knockdown of Cno, a subset of AJs become destabilized during germband elongation (Manning et al., 2019; Perez-Vale et al., 2021; Sawyer et al., 2011), particularly TCJs and those along aligned AP borders, which are known or suspected to be under elevated tension (Fig 2G, arrows). We thus examined the frequency of AJ defects in *cnoΔProxIDR* mutants at stages 7 and 8, using staining for Arm. We included places with small or large gaps where three or more cells came together and places where AJ protein accumulation was broadened, and scored gaps blinded in wildtype and *cnoΔProxIDR* mutants. Mutants had a dramatically elevated number of gaps at multicellular junctions—a mean of 5.5 gaps per field in wildtype and 38 gaps per field in *cnoΔProxIDR* mutants (Fig 2H vs 2I; quantified in 2J; the one outlier with near normal gap numbers among the fourteen *cnoΔProxIDR* mutants might have been a zygotically rescued embryo). However, there was a qualitative difference between *cnoΔProxIDR* mutants and strong *cno* mutants—instead of the many broad gaps at AJs seen in strong mutants (Fig 2G, arrows), we primarily saw smaller gaps in *cnoΔProxIDR* mutants (Fig 2I, I’ arrows).

During stages 9-10 cells in the lateral and ventral ectoderm experience two challenges to AJs, those posed by mitosis and those posed by invagination of ∼30% of cells as neural stem cells. These challenges make the ventral epidermis particularly susceptible to reduced cell adhesion. In wildtype most cells are columnar, as they rapidly return to this shape after rounding up for division (Fig 2K). In contrast, the recovery of columnar cell shape after division is delayed in strong *cno* mutants, ultimately leading to penetrant problems with ventral epidermal integrity (Manning et al., 2019; Perez-Vale et al., 2021). At stage 9, most *cnoΔProxIDR* mutants appeared relatively normal, aside from continuing issues with failure to close the ventral furrow. However, at stages 10-11 many embryos had ventral cells that failed to return to columnar shape after division. These defects were variable in severity: ∼38% presented with only mild defects (10/26 embryos), 35% had more widespread failure of columnar cell architecture (9/26 embryos; Fig 2L, arrows), and 27% had more severely disrupted ventral epidermis (7/26 embryos; Fig 2M, N, arrows). After germband retraction during stages 13-14, wildtype embryos complete the final two events of epidermal morphogenesis, head involution and dorsal closure (Fig 2O, yellow and cyan arrows, respectively). Consistent with the observed cuticle defects, a subset of *cnoΔProxIDR* mutants failed to close dorsally (Fig 2R, cyan arrows), head involution was often defective (Fig 2R, S, yellow arrows), and leading-edge cell shapes were hyper-constricted or hyper-extended (Fig 2P vs Q), as we observed in strong *cno* mutants (Manning et al., 2019). However, few embryos had ventral epidermal holes, consistent with the cuticles, another way in which *cnoΔProxIDR* mutants are less severe than the null mutant.

Our final analysis of proximal IDR function was to examine its role in AJ substructure. Cadherin/catenin complexes are not continuous along the membrane but assemble into large puncta containing hundreds of proteins (McGill et al., 2009). Enhanced resolution imaging using the Airyscan module revealed that this punctate array of Arm is maintained during germband extension (Fig 2T, yellow arrows; (Schmidt et al., 2023). Intriguingly, Cno’s C-terminus is also enriched in puncta, but many puncta are differentially enriched for Arm vs Cno (Fig 2T, yellow vs cyan arrows; (Schmidt et al., 2023). When we examined *cnoΔProxIDR* mutants we saw no obvious changes in the punctate nature of Arm localization along the AJ (Fig 2U, yellow arrows). Loss of Polychaetoid/ZO1 similarly does not alter Arm’s punctate localization (Schmidt et al., 2023). While CnoΔProxIDR localization to AJs was substantially reduced, if we enhanced the junctional signal, it appeared to localize to a broader zone than is occupied by Arm puncta (Fig 2U, yellow vs cyan arrows). Thus, in strong contrast to our earlier mutants in which folded domains were deleted, the proximal IDR is critical for Cno AJ localization, Cno function in AJ stabilization, and many aspects of morphogenesis. However, subtle differences with the *cno* null mutant phenotype suggest CnoΔProxIDR retains some residual function.

### Replacing Cno’s proximal IDR with Afadin’s restores AJ localization and viability but does not provide full function

While the five folded protein domains of Cno and Afadin are well conserved (42-74% sequence identity; (Gurley et al., 2023)) and the C-terminal FAB has clear blocks of conservation (35% identity), the proximal IDR is highly divergent between flies and mammals (Gurley et al., 2023). We thus began with the hypothesis that Afadin’s proximal IDR would not functionally replace that of Cno. To test this, we inserted into the *cno* locus a mutant with Cno’s proximal IDR replaced Afadin’s (*cnoAfadinIDR*; Fig 3A). As an N-terminal boundary for the replacement we began 30 amino acids downstream of the PDZ domain, as the preceding region was well conserved among different insects and different vertebrates and may be a PDZ domain extension. For the C-terminal boundary we ended the deletion 10 amino acids N-terminal to our earlier FAB deletion. For cloning, we engineered restriction sites into these positions encoding short linker peptides (Methods and Fig S1). As a control, we re-inserted the Cno IDR coding sequence into the same location (*cnoCanoeIDR*; Fig 3A). We verified that CnoCanoeIDR accumulated at near-wildtype levels, while CnoAfadin IDR accumulated at somewhat lower levels (∼60%; Fig S2C, D).

**Fig 3.**
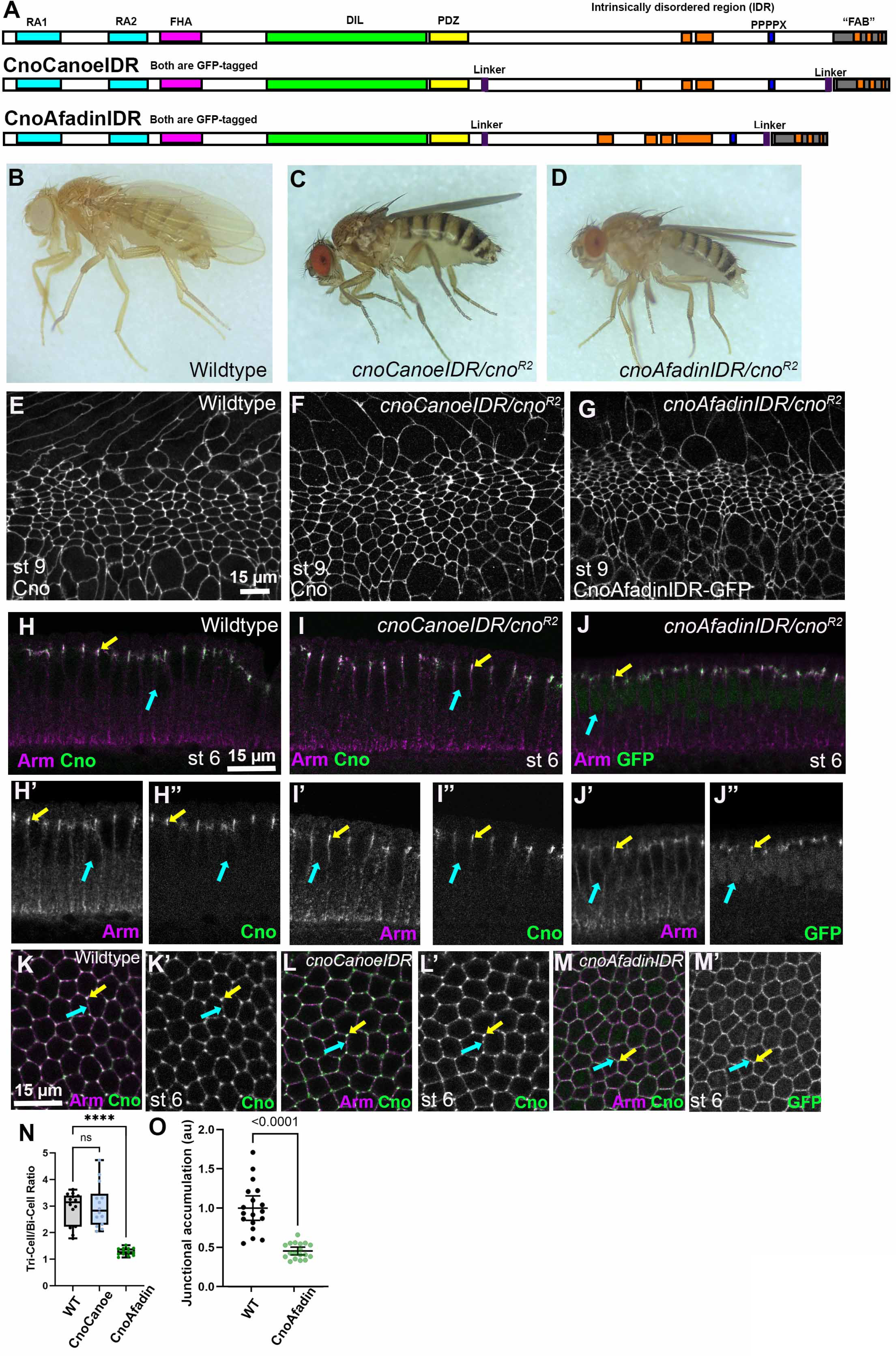
Afadin’s IDR can restore viability and fertility and largely but not complete restore protein localization. **(A)**. Drosophila Cno and the IDR replacement mutants. **(B-D).** Maternal/zygotic mutant adults of the indicated genotype. **(E-M).** Embryos, anterior left, antigens and stages indicated. **(E-G).** Stage 9. Both mutant proteins localize to AJs. **(H-J).** Cross-sections, apical up, stage 6. Yellow arrows=junctional localization. Cyan arrows=nuclei. **(K-M).** Tricellular junction enrichment. Yellow arrows=TCJs, cyan arrows=bicellular junctions. **(N).** Quantification of Tricellular junction enrichment**. (O).** Relative accumulation at AJs.

We then crossed each mutant to our standard null allele, *cno^R2^*. To our surprise, *cnoAfadinIDR/cnoR^2^* flies were viable to adulthood, as were the *cnoCanoeIDR/cno^R2^* control (Fig 3C, D). However, while *cnoCanoeIDR/cno^R2^* adults appeared at near-Mendelian ratios (26% versus 33% expected (n=827)), *cnoAfadinIDR/cno^R2^*adults were less viable (16% versus 33% expected (n=1167)). Both produced viable progeny surviving to adulthood, confirming that maternal-zygotic mutants are viable. Both CnoCanoeIDR (Fig 3F) and CnoAfadinIDR (Fig 3G) proteins localized to AJs, with enrichment at the apical end of the lateral interface, like wildtype Cno (Fig 3I, J vs H, yellow arrows). However, replacing Cno’s proximal IDR with that of Afadin did not fully eliminate the nuclear localization seen in CnoΔProxIDR (Fig 3J, cyan arrow). We also assessed TCJ enrichment. While CnoCanoeIDR enrichment at TCJs matched that of wildtype Cno (Fig 3L vs K, arrows; quantified in N), CnoAfadinIDR exhibited substantially reduced TCJ enrichment (Fig 3M vs K, arrows; quantified in N). Finally, we examined the overall levels of AJ localization, comparing the progeny of *cnoAfadinIDR/cno^R2^* parents to wildtype CanoeGFP on the same slide, using the GFP signal. CnoAfadinIDR localization to AJs was reduced about 2-fold (Fig 3O). Thus, replacing Cno’s proximal IDR with that of Afadin restores AJ localization and adult viability, but does not restore enrichment at TCJs under tension.

In our previous work we developed a sensitized assay to assess function of viable alleles (Perez-Vale et al., 2021). To do so, we reduced the maternal dose of the mutant protein by making mothers transheterozygous for our null allele, *cno^R2^*, and then crossed transheterozygotes to one another. When we did this assay with the wildtype allele, crossing *+/cno^R2^* males and females, 25% of the progeny die: those that are *cno^R2^/cno^R2^*. Because of the strong maternal contribution, the dead embryos only have mild defects in morphogenesis, as assessed by examining cuticles, primarily defects in head involution (Fig 4A; (Gurley et al., 2023). Dorsal closure and epidermal integrity, which are disrupted in maternal/zygotic null mutants, remain largely unaffected.

**Fig 4.**
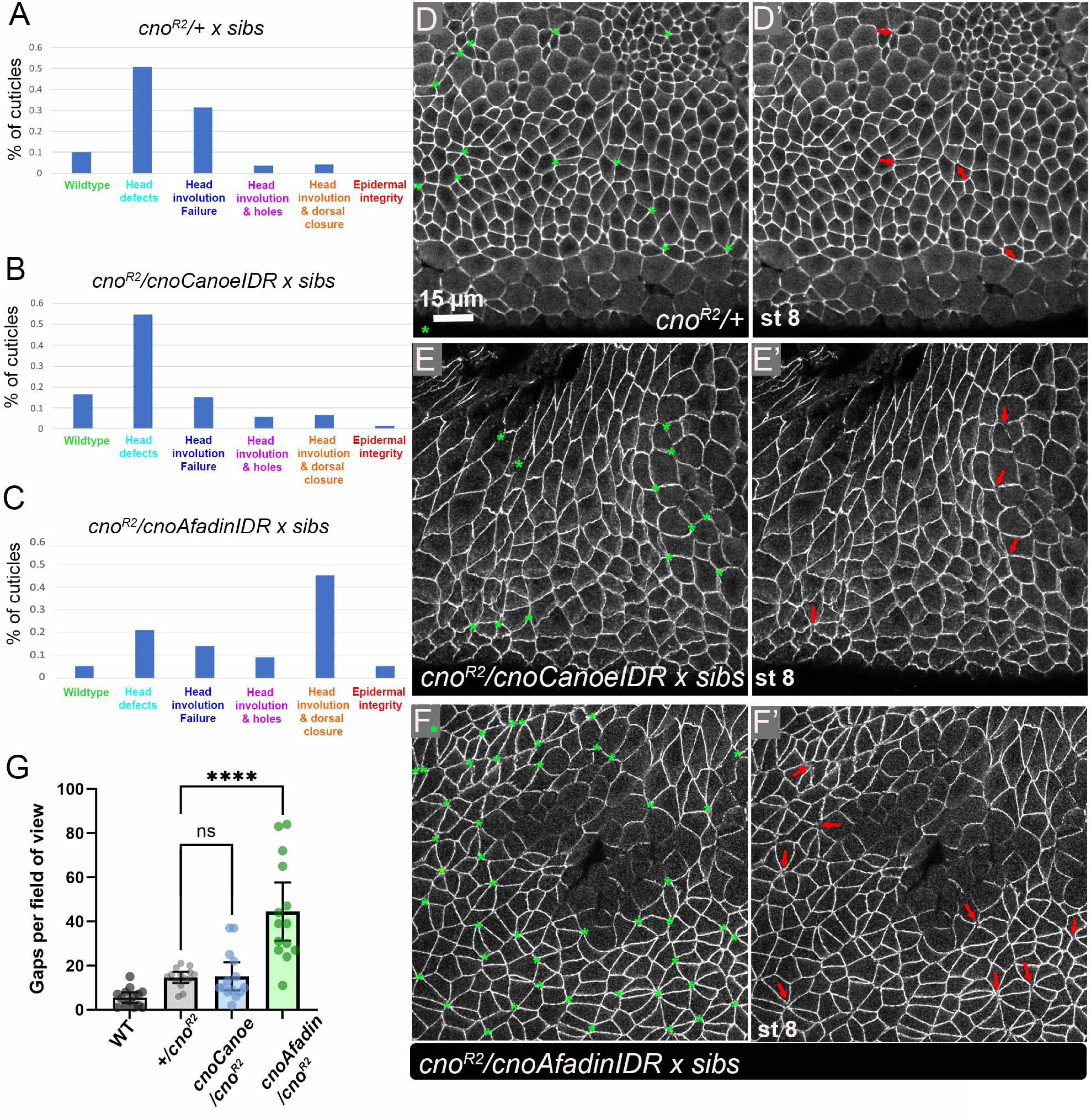
Sensitized assays reveal that *cnoAfadinIDR* does not have fully wildtype function. **(A-C)**. Distribution of cuticle phenotypes in different mutants, using cuticle examples in Fig 1. *cno^R2^/+* data is from (Gurley et al., 2023). **(D-F).** Junctional gaps, Stage 8. Asterisks=all junctional gaps/disruptions. Arrows=selected gaps. **(G).** Gap quantification.

We tested the function of CnoCanoeIDR and CnoAfadinIDR using this assay. When we crossed *cnoCanoeIDR /cno^R2^* males and females, we saw 42% lethality (n=1317), above the 25% expected if it provided full function. However, the cuticles of dead embryos only had mild defects in head involution, comparable to those we saw when crossing *+/cno^R2^* males and females (Fig 4B vs A). Thus, this allele provides substantial but not full function, perhaps due to effects of the small disruptions generated in the IDR at the two ends of our replacement allele. CnoAfadinIDR provided substantially less function. When we crossed *cnoAfadinIDR /cno^R2^* males and females, we saw 64% lethality (n=675), suggesting many *cnoAfadinIDR /cno^R2^* embryos die. Further, the cuticle phenotypes were substantially enhanced, with 45% of embryos exhibiting failure of both head involution and dorsal closure (Fig 4C vs A). These data suggest that the Afadin IDR provides enough function for viability, but this function is not fully wildtype. To further this analysis, we examined the ability of each IDR replacement to stabilize AJs under tension, using as a control the progeny of

*+/cno^R2^* parents. These control embryos had a slightly elevated number of AJ gaps relative to wildtype. Progeny of *cnoCanoeIDR /cno^R2^* parents also had slightly elevated numbers of AJ gaps, similar to our controls (Fig 4E vs D, green asterisks; quantified in G). In contrast, progeny of *cnoAfadinIDR /cno^R2^* parents had highly elevated junctional gap number (Fig 4F vs D; quantified in G). It is important to note there are three genotypes among the progeny (*mutant/mutant, mutant/cno^R2^,* and *cno^R2^/cno^R2^)*; we think this likely contributes to the variability in gap number between embryos. Finally, we scored embryos for defects in ventral furrow invagination. 6% of the progeny of +/*cno^R2^* parents have mild defects in this process (McParland et al., 2024a). Progeny of *cnoCanoeIDR /cno^R2^* parents displayed a slightly elevated defect frequency (11%), with most defects being minor, whereas progeny of *cnoAfadinIDR /cno^R2^* parents exhibited a substantially higher incidence of ventral furrow defects (49%; Table S1). Taken together, these data reveal that Afadin’s IDR can function in the Cno context, restoring AJ localization and sufficient protein function to convert an embryonic lethality to adult viability and fertility. However, examination of enrichment at AJs under tension and sensitized functional assays revealed that it does not restore full wildtype function.

### ΑlphaFold predicts conserved helical regions in the IDR, some of which have defined functions in Afadin

Many IDRs contain short motifs serving as binding sites for other proteins. While reciprocal BLAST searching did not reveal obvious conserved sequences in the proximal IDR, ΑlphaFold provided an alternate approach to look for elements conserved at the structural level. Afadin’s proximal IDR contains four predicted α-helices (Fig 3A, orange boxes(Gurley et al., 2023). The most N-terminal helix precisely overlaps the known binding site for the M-domain of αcat (Maruo et al., 2018). Intriguingly, this helix is clearly absent from the Cno IDR. Cno only has three predicted α-helices, a nine amino acid glutamine-rich helix and two longer helices.

When we used the sequence of the two longer helices to BLAST the human proteome, it revealed similarity to two of the Afadin helices, with 33% sequence identity over 72 amino acids (Gurley et al., 2023). Our interest in this conserved region was substantially heightened by recent cryo-EM studies from the Alushin lab using mammalian proteins. This conserved region of Afadin folds as a forked pair of α-helices and forms a ternary complex with the actin-binding domain of αcat and an actin filament, enhancing their interaction (Gong et al., 2025). We thus hypothesized this region would play an important role in Cno function.

### Neither the conserved α-helices nor the IDR’s N- or C-terminal regions are individually essential for viability, but the N- and C-terminal regions are required for full function

We used the two conserved α-helices to subdivide the IDR into three regions: the helices themselves and the regions N-terminal or C-terminal to them. To evaluate the extent to which these regions contribute to Canoe function, we created a set of mutants (Fig 5A) lacking the conserved helices (cnoIDRΔH), the region N-terminal to them (cnoIDRΔN; ending 14 amino acids before the predicted helices), or the region C-terminal to them (cnoIDRΔC; beginning 9 amino acids after the predicted helices; details in Methods/Fig S1). All three accumulated at roughly wildtype levels (Fig S2E, F), Our initial hypothesis was that the conserved helices would be essential. We first examined whether these IDR regions were essential for adult viability, by crossing each mutant to our null allele, *cno^R2^*. Strikingly, cnoIDR*ΔH/cno^R2^, cnoIDRΔN/cno^R2^*, and *cnoIDRΔC/cno^R2^* flies were all viable to adulthood and produced viable progeny that survived to adulthood. After outcrossing we obtained homozygous mutant lines of each (Fig 4B-E). However, while *cnoIDRΔH/cno^R2^* progeny were obtained at Mendelian ratios (33% versus the expected 33% (Balancer homozygotes die; n= 602), both *cnoIDRΔN/cno^R2^* and *cnoIDRΔC/cno^R2^*flies appeared at less than Mendelian ratios (19% for *cnoIDRΔN/cno^R2^*, n= 514; 12% for *cnoIDRΔC/cno^R2^*, n= 523). Thus, none of these three IDR regions is essential for viability or fertility, but the reduced viability suggested two of the regions contribute to full function.

**Fig 5.**
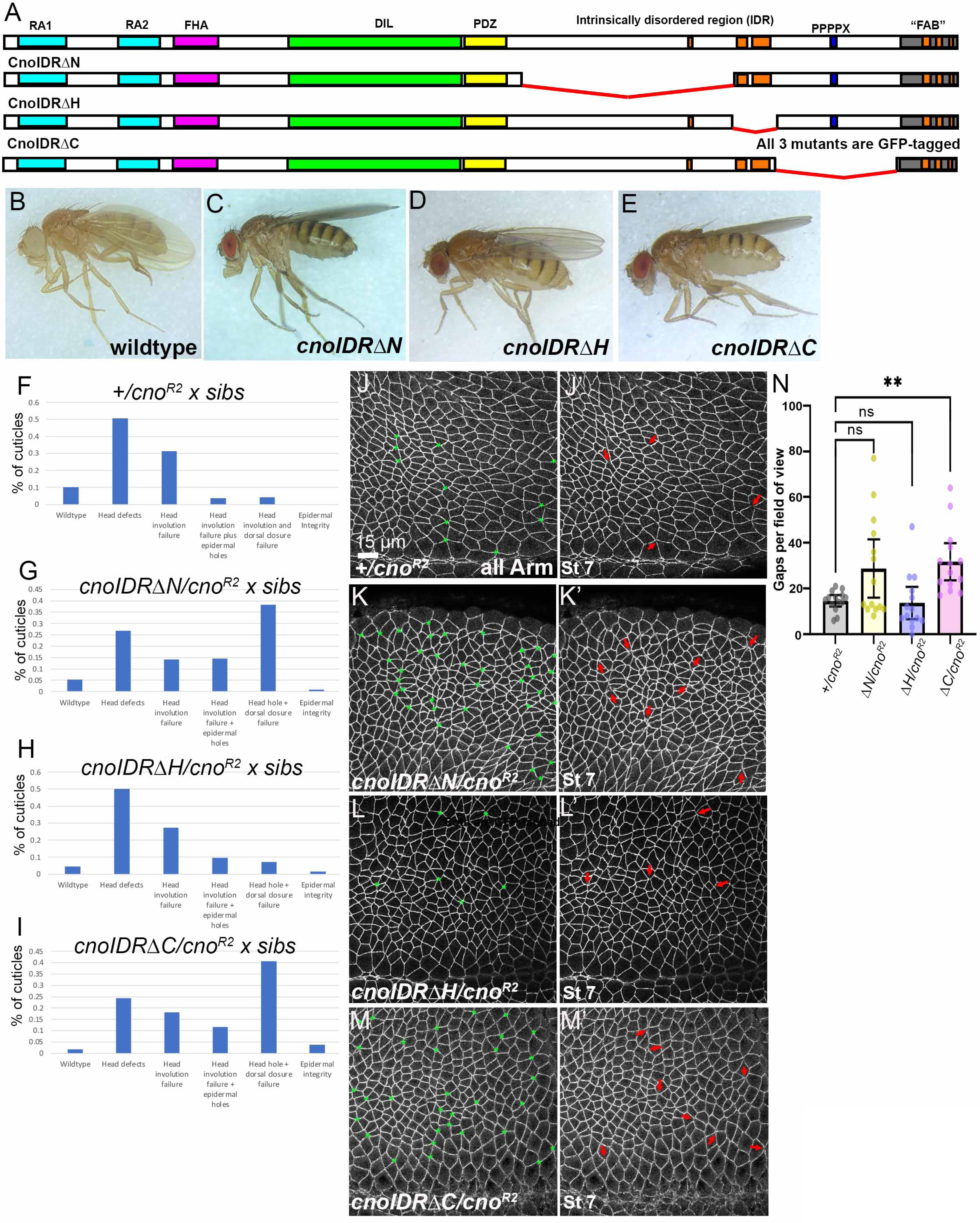
Neither the conserved α-helices nor the regions of the IDR N- or C-terminal are individually essential for viability, but the N- and C-terminal regions of the IDR are required for full wildtype function. **(A).** Drosophila Cno and the IDR deletion mutants. **(B-D).** Maternal/zygotic adults of the indicated genotype. **(F-I).** Distribution of cuticle phenotypes in different mutants, using cuticle examples in Fig 1. *cno^R2^/+* data is from Gurley et al. (Gurley et al., 2023). **(J-M).** Junctional gaps, Stage 7. Asterisks=all junctional gaps/disruptions. Arrows=selected gaps. **(N).** Gap quantification.

We used our sensitized assay to test this further, crossing males and females transheterozygous for each allele and our null allele, *cno^R2^*. While the conserved helices were not essential for viability, we still thought they would be the most important region of the proximal IDR. However, when we crossed *cnoIDRΔH /cno^R2^* males and females, we saw 29% lethality (n=785), only slightly above the 25% expected if it provided full function.

Cuticle phenotypes of the dead embryos were only slightly more severe than those of *cno^R2^* zygotic mutants, with 81% of embryos exhibiting defects restricted to head involution (Fig 5H vs F). Thus CnoIDRΔH provides nearly wildtype function. In contrast, the other two mutants provided less function. Crossing *cnoIDRΔN /cno^R2^* males and females led to 34% lethality (n=690), whereas the same cross with *cnoIDRΔC /cno^R2^* males and females resulted in 53% lethality (n=866). In both cases, cuticle phenotypes were substantially enhanced, with 38% (Fig 5G) or 41% (Fig 5I) of the embryos having failure of both dorsal closure and head involution.

Deleting the full IDR led to frequent gaps at multicellular junctions (Fig 2I). To determine which IDR regions are required to stabilize AJs under tension, we examined AJ gap frequency in each mutant, crossing *mutant/cno^R2^* males and females, using progeny of *+ /cno^R2^* parents as a control. *cnoIDRΔH/cno^R2^* mutants had a slightly elevated frequency of gaps, similar to our +*/cno^R2^* control (Fig 5L vs J, quantified in N) or to *cnoCanoeIDR/cno^R2^* mutants. Mean gap frequency trended higher in *cnoIDRΔN/cno^R2^* mutants but did not reach statistical significance (Fig 5K vs J, quantified in N). Gap frequency was substantially elevated in *cnoIDRΔC/cno^R2^* mutants (Fig 5M vs J, quantified in N). Finally, we scored the fidelity of ventral furrow invagination (Table S1), using as a baseline our previous analysis of progeny of +*/cno^R2^* parents (6% defects; (McParland et al., 2024a)). Progeny of *cnoIDRΔH/cnoR2* parents exhibited a slightly elevated frequency of defects (11%, all mild; Table S1)—this was somewhat more elevated in progeny of *cnoIDRΔN/cno^R2^* parents (19%; Table S1). In contrast, both the frequency and severity of defects was substantially elevated in progeny of *cnoIDRΔC/cno^R2^* parents (52%; Table S1). Together these data reveal that none of the individual IDR regions are essential to restore viability and fertility. While conserved helices appear largely dispensable for function, the N- and C-terminal IDR regions are required for full wildtype function, with the C-terminal region playing a more important role.

### The IDR’s C terminal region is important for nuclear exclusion and together with the N-terminal region contributes to TCJ enrichment

CnoΔProxIDR mutant protein exhibited reduced localization to AJs, loss of TCJ enrichment and surprising mis-localization to nuclei (Fig 1). We thus examined whether any individual proximal IDR region was important for AJ localization. All three mutant proteins clearly localized to AJs at embryonic stage 9 (Fig 6A vs B-D). In cross section, both CnoIDRΔN and CnoIDRΔH proteins were strongly enriched at apical AJs (Fig 6F,G vs E, yellow arrows). We next examined enrichment of each at TCJs, where wildtype Cno is enriched in response to mechanical tension (Fig 6I, yellow vs cyan arrows). CnoIDRΔH enrichment at TCJs relative to bicellular junctions was unchanged from wildtype (Fig 6K, yellow vs cyan arrows, quantified in M). However, CnoIDRΔN enrichment at TCJs was reduced, though the protein retained ∼2-fold enrichment (Fig 6J, yellow vs cyan arrows, quantified in M).

**Fig 6.**
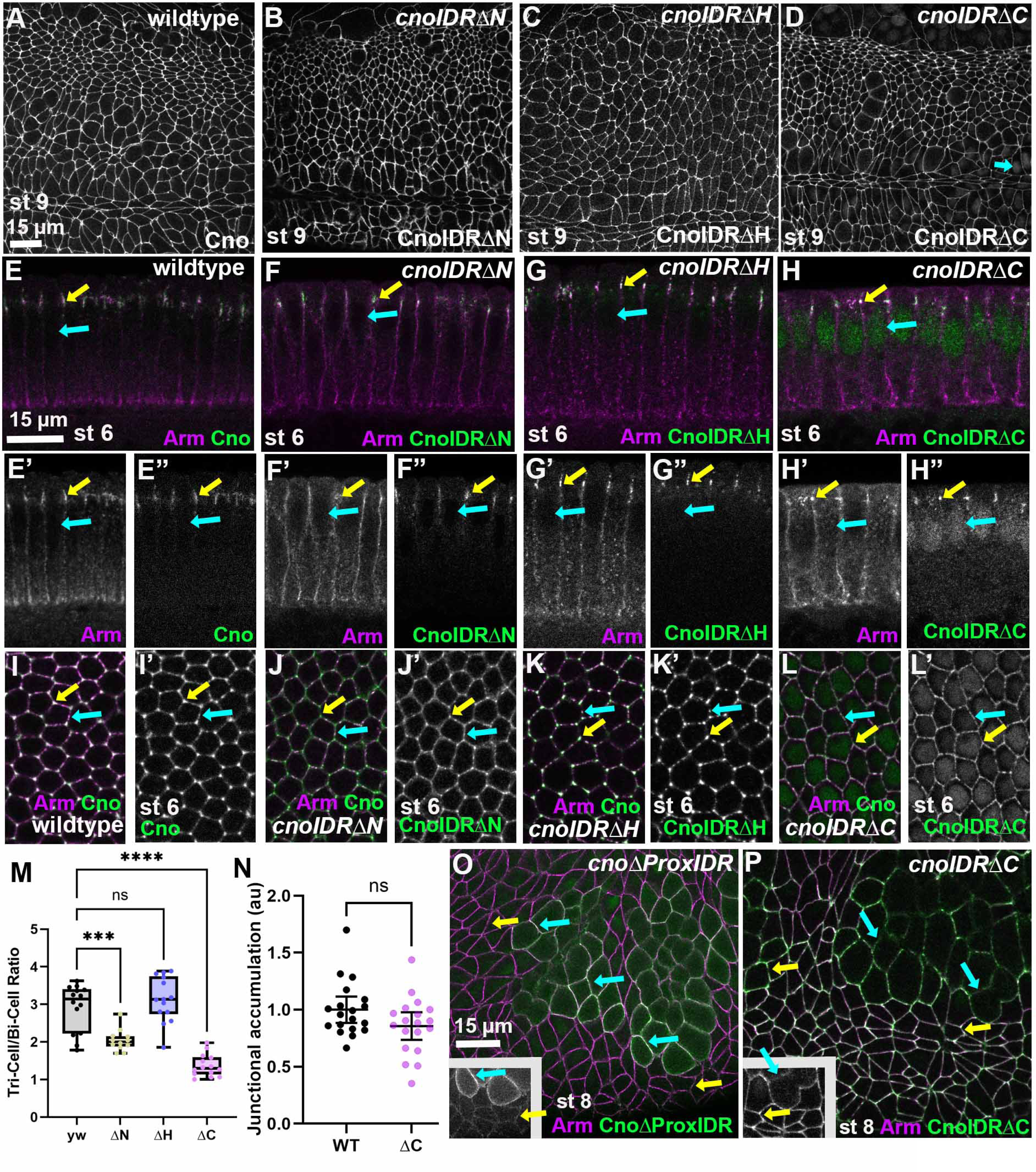
The C terminal region of the IDR is important for nuclear exclusion and with the N-terminal region contributes to TCJ enrichment. **(A-D).** Stage 9. All mutant proteins localize to AJs, although CnoIDRΔC also localizes to nuclei (D, arrow). **(E-H).** Cross-sections, apical up, stage 6. Yellow arrows=junctional localization. Cyan arrows=nuclei. **(I-M).** Tricellular junction enrichment. Yellow arrows=TCJs, cyan arrows=bicellular junctions. **(N).** Relative accumulation of CnoIDRΔC at AJs. **(O, P).** Stage 8. Cyan arrows=cells rounded up for mitosis. Yellow arrows=nondividing cells.

In contrast, CnoIDRΔC protein localization was substantially altered. While it accumulated strongly at apical AJs (Fig 6D, H yellow arrow), cross-sections revealed it also accumulated in nuclei (Fig 6H, cyan arrow), resembling CnoΔProxIDR in this regard. TCJ enrichment of CnoIDRΔC protein was essentially eliminated (Fig 6L yellow vs cyan arrows, quantified in M), but levels of CnoIDRΔC protein were not significantly different from wildtype (Fig 6N). While CnoIDRΔC and CnoΔProxIDR exhibited similar accumulation in nuclei, other aspects localization were not shared. CnoΔProxIDR accumulation in AJs of non-dividing cells was substantially reduced relative to wildtype (Fig 1S; 6O yellow arrows), whereas CnoIDRΔC protein accumulated in AJs of non-dividing cells at essentially wildtype levels (Fig 6P, yellow arrows). In addition, as cells round up for mitosis and nuclei disassemble, CnoΔProxIDR protein is released into the cytoplasm and accumulates at high levels at the mitotic cell cortex (Fig 1S, 6O, cyan arrows). In contrast, CnoIDRΔC protein resembles wildtype Cno, with somewhat reduced levels in mitotic cells (Fig 6P, cyan arrows).

### The conserved helices are not sufficient for Cno localization or function

Thus, both the N- and C-terminal regions of the proximal IDR are dispensable for viability. Since the two helices are the only long conserved sequence in the proximal IDR and since, in Afadin, they have a clear role in stabilizing the interaction of αcat with actin (Gong et al., 2025), we tested the hypothesis that they are sufficient for IDR function. To do so, we created a mutant retaining only the conserved helices in the proximal IDR (*cnoIDRHelixOnly* (Fig 7A), thus combining the regions deleted in CnoIDRΔN and CnoIDRΔC (Fig 5A). CnoIDRHelixOnly accumulates at near wildtype levels (Fig S2A, B). *cnoIDRHelixOnly* was not viable over our null allele, *cno^R2^* (0/558 progeny), thus ruling out the idea that the helices are sufficient for function. However, unlike *cno^R2^*, but similar to *cnoΔProxIDR*, *cnoIDRHelixOnly* was not zygotically embryonic lethal (2% lethality, n=720), suggesting it might retain a small amount of function.

**Fig 7.**
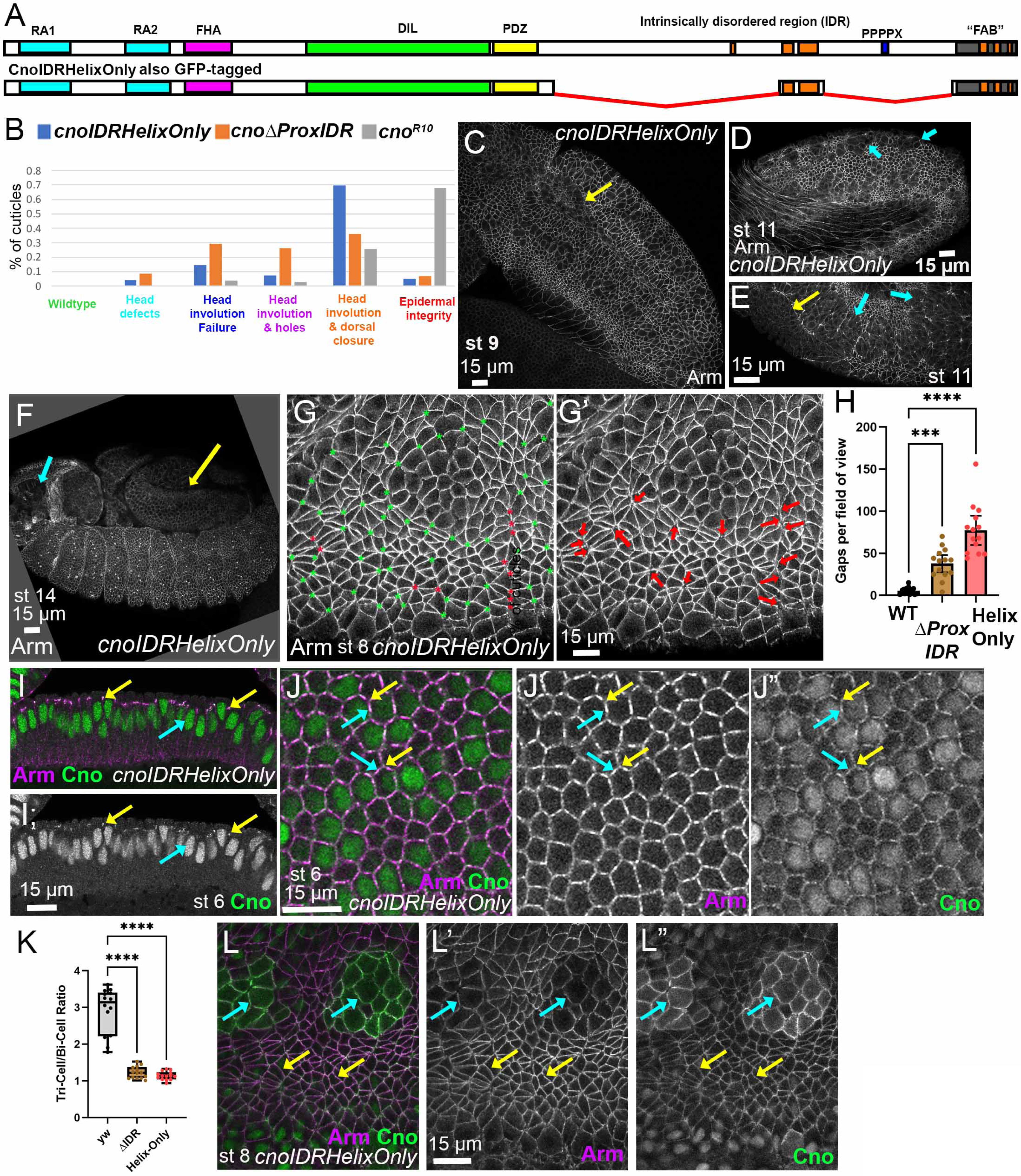
The conserved helices are not sufficient for Cno localization or function. **(A)**. Drosophila Cno and CnoHelixOnly mutant. **(B).** Distribution of cuticle phenotypes in different mutants. *cno^R10^/+* data is from (Gurley et al., 2023). **(C-F).** *cnoHelixOnly* embryos, anterior left, antigens indicated. **(C, E).** Defects in ventral furrow invagination (yellow arrows). **(D, E).** Defects in resuming columnar cell architecture (cyan arrows). **(F).** Failure of dorsal closure (yellow arrow) and head involution (cyan arrow). **(G).** Junctional gaps, Stage 8. Asterisks=all junctional gaps/disruptions. Arrows=selected gaps. **(H).** Gap quantification. **(I).** Cross-sections, apical up, stage 6. Yellow arrows=junctional localization. Cyan arrows=nuclei. **(J, K).** TCJ enrichment. Yellow arrows=TCJs, cyan arrows=bicellular junctions. **(L).** Stage 8. Cyan arrows=cells rounded up for mitosis. Yellow arrows=nondividing cells.

To evaluate CnoIDRHelixOnly function, we generated maternal-zygotic mutants. We observed 55% lethality (n=727), consistent with fully penetrant lethality of maternal-zygotic mutants and substantial zygotic rescue. We next assessed function in morphogenesis by examining cuticle phenotypes. Strikingly, *cnoIDRHelixOnly* maternal-zygotic mutants had strong defects in morphogenesis that were, on average, more severe than those of mutants lacking the entire IDR (Fig 7B). 70% of embryos exhibited failure of both head involution and dorsal closure, in contrast to only 31% of *cnoΔProxIDR* mutants. However, the defects in morphogenesis were not as severe as maternal-zygotic null mutants, with only 5% having additional defects in epidermal integrity, unlike the 68% in null mutants (Fig 7B). Next, we looked at embryos as development proceeded. *cnoIDRHelixOnly* mutants had moderate to strong defects in ventral furrow invagination (Fig 7C, E, yellow arrows; 78% defective, n=82; Table S1). Moreover, *cnoIDRHelixOnly* mutants also exhibited delays in resuming columnar cell architecture after later cell divisions (Fig 7D, E, cyan arrows) and had defects in both head involution and dorsal closure (Fig 7F, cyan vs yellow arrows), like strong *cno* mutants. To look more closely at underlying AJ defects, we examined AJ gaps. *cnoIDRHelixOnly* maternal-zygotic mutants had an exceptional number of AJ gaps, with many if not most multicellular junctions affected. These were more frequent than those seen in *cnoΔProxIDR* (Fig 7G, quantified in H) and included broader gaps along aligned AP borders (Fig 7G, red asterisks).

CnoIDRHelixOnly protein accumulated in nuclei, with reduced localization to AJs (Fig 7I-J). TCJ enrichment of the mutant protein was virtually eliminated (Fig 7J, yellow arrows, quantified in K). During germband extension, AJ localization in non-mitotic cells remained relatively weak (Fig L, yellow arrows), but, as we saw for CnoΔProxIDR protein, in mitotic cells CnoIDRHelixOnly protein accumulated at high levels at the cortex (Fig 7L, cyan arrows). Thus, the conserved helices are not sufficient for Cno localization or for Cno function, and some phenotypes of *cnoIDRHelixOnly* mutants are more severe than those of *cnoΔProxIDR*.

### The IDR also plays an important role in Cno function during post-embryonic eye development

Cno also plays important roles in postembryonic development. For example, in the developing eye a field of undifferentiated imaginal disc cells re-organizes through orchestrated cell shape changes into an array of identical ommatidia. Each has a defined number of cone cells, primary (1°), secondary (2°), and tertiary (3°) cells, and bristles, arranged in precise relationships to one another (Fig 8A; (Johnson, 2021). AJ proteins and their regulators play important roles in the eye (Johnson, 2021). Complete loss of Cno dramatic disrupts the epithelium (Walther et al., 2018), while milder mutants have less severe defects (e.g. (Matsuo et al., 1997; Matsuo et al., 1999; Miyamoto et al., 1995; Takahashi et al., 1998). Cno’s regulator Rap1 also plays important roles (Yost et al., 2023). Even *cno* alleles that are adult viable and fertile, like *cnoΔPDZ,* exhibit defects in ommatidial cell number and arrangement (McParland et al., 2024a; McParland et al., 2024b; Perez-Vale et al., 2021), providing another tissue in which to assess function.

**Fig 8.**
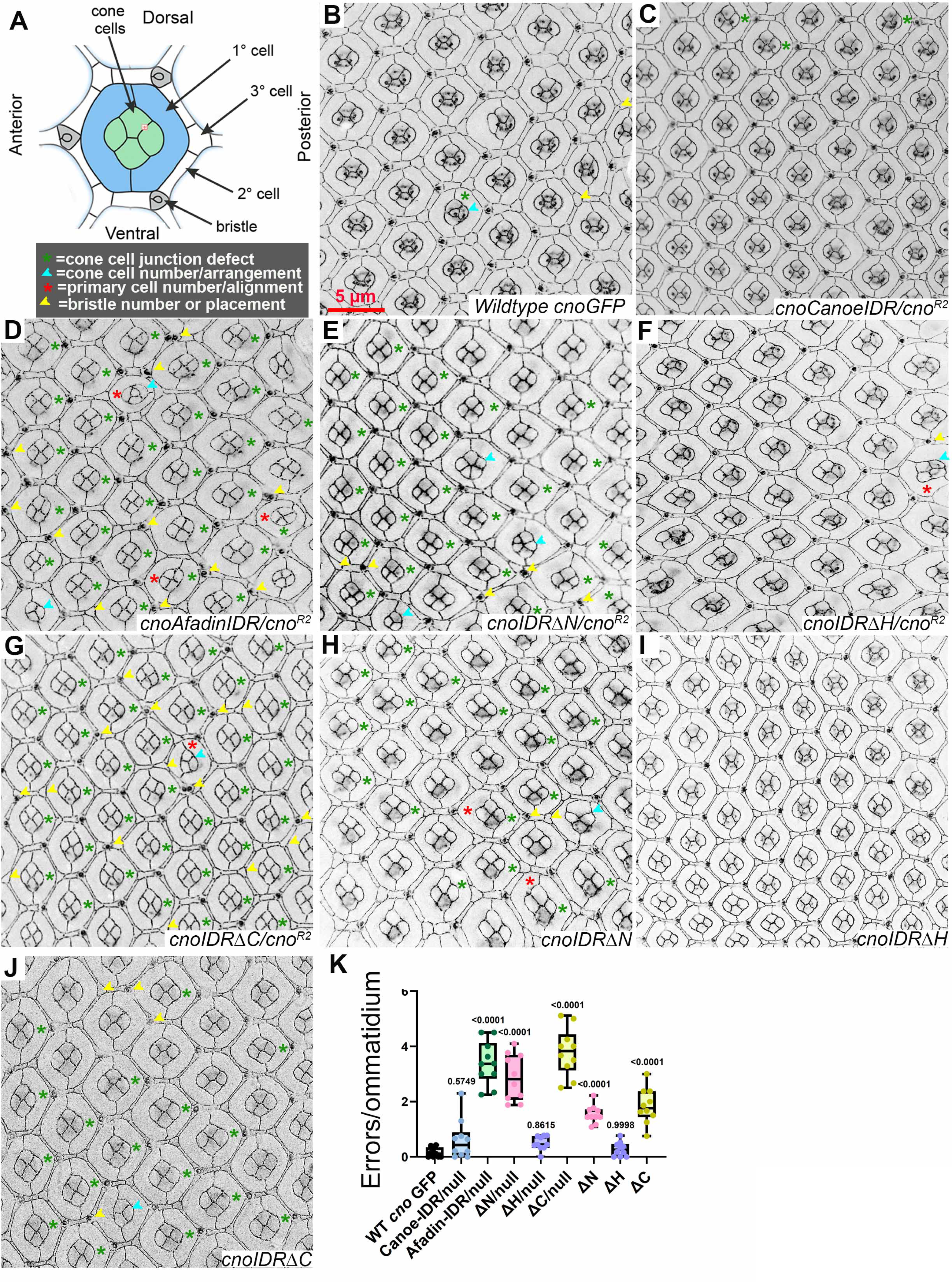
Multiple regions of the IDR are important for Cno function in the developing eye. **(A)**. Diagram of the complex cell arrangement in a wildtype ommatidium with a key to defects observed in mutants. **(B-J).** Representative pupal eyes from each genotype, highlighted to illustrate defects observed. **(K).** Quantification of mean number of defects per ommatidium with statistical analysis of difference from wildtype CnoGFP.

We thus generated transheterozygous animals with each of our new alleles over the *cno^R2^* null allele, using our GFP-tagged wildtype Cno as a control. Our previous work revealed that +/*cno^R2^*animals do not have significant numbers of defects (McParland et al., 2024b). Following dissection at 40 hours after puparium formation, eye discs were stained and imaged for Ecad and N-cadherin and additionally imaged in the GFP-channel to visualize mutant Cno proteins. For each eye disc, we scored defects in the stereotyped cell arrangement using an established scoring system (Johnson and Cagan, 2009), scoring defects in the number and arrangement of cone cells, 1* cells, 3* cells and bristles (Table S2). We scored 10 eye discs with 77-115 total ommatidia per genotype.

Wildtype eyes have occasional defects: ∼0.5 defects per ommatidium. Our wildtype GFP-tagged *cno* mutant has a defect frequency similar to wildtype (Perez-Vale et al., 2021). We replicated that here (Fig 8B, K; 0.52 defects/ommatidium), and it was our baseline control. For *cnoCanoeIDR* and *cnoAfadinIDR* we scored animals heterozygous with the null allele *cno^R2^*, as we never obtained a homozygous *cnoAfadinIDR* line. *cnoCanoeIDR/cno^R2^* mutants had a defect number that was not significantly elevated (Fig 8C, K; 0.61 defects/ommatidium). In contrast, *cnoAfadinIDR/cno^R2^* mutants had a substantially elevated number of defects (Fig 8D, K; 3.4 defects/ommatidium). We next examined mutants carrying smaller deletions in the IDR. *cnoIDRΔH/cno^R2^* mutants had defect frequency similar to wildtype (Fig 8E, K; 0.50 defects/ommatidium). In contrast, both *cnoIDRΔN /cno^R2^* mutants (Fig 8E, K) and *cnoIDRΔC/cno^R2^* mutants (Fig 8G, K) had substantially elevated numbers of defects (2.9 and 3.8 defects/ommatidium, respectively). With these mutants we also could look at homozygotes and saw similar, though weaker, effects (Fig, 8 H-K; *cnoIDRΔN*=1.5 defects/ommatidium; *cnoIDRΔH*=0.50 defects/ommatidium; *cnoIDRΔC*=1.8 defects/ommatidium). These results precisely paralleled each of our sensitized assays in embryos, with CnoCanoeIDR and CnoIDRΔH retaining near wildtype function, CnoIDRΔN modestly reduced in function and CnoAfadinIDR and CnoIDRΔC providing significantly less than wildtype levels of function.

### Both the conserved helices and the FAB can bind F-actin

The surprising robustness of the IDR raised significant questions. No individual region of the proximal IDR+FAB was essential for viability and all mutants retained substantial function. We were particularly surprised at the dispensability of the conserved helices, as they are the only region of Afadin known to bind F-actin directly (Carminati et al., 2016; Gong et al., 2025). In contrast, while the C-terminal IDR is traditionally referred to as the FAB (most recently defined as the C-terminal 77 amino acids; (Sakakibara et al., 2020), no one ever directly assessed whether it can bind F-actin.

These data prompted us to directly assess the ability of different regions of Cno’s proximal IDR and FAB to bind actin. We utilized an F-actin co-sedimentation assay, in which actin monomers are polymerized in vitro into filaments. Following centrifugation, the large actin filaments pellet (Fig 9A). One can thus assess whether other proteins bind F-actin by evaluating whether they co-sediment with the filaments. We used the actin-binding domain of αcat as a positive control. When centrifuged alone, it remained in the supernatant, but 70% pelleted when mixed with F-actin (Fig 9A, E). The only previous assessment of Cno’s ability to bind F-actin used a long construct extending from just before the conserved helices to the end of the Cno (Sawyer et al., 2009), similar to the construct originally used to define Afadin’s ability to bind F-actin (Mandai et al., 1997). We first replicated our earlier work with a fragment of Cno extending from 6 amino acids N-terminal to the conserved helices to Cno’s C-terminus (referred to as H+C+FAB; Fig S3). This was tagged at its N-terminus with GST for purification and mixed with F-actin. 32% co-sedimented with F-actin (Fig 9B, F). We next examined whether adding αcat’s actin binding domain enhanced interaction of H+C+FAB with F-actin. The increase did not reach statistical significance (Fig 9B, F). When we purified H+C+FAB, we obtained both the full-length fusion protein (Fig 9B, blue arrowhead) and several C-terminal degradation products (Fig 9B, red arrowhead). Intriguingly, the truncated products co-sedimented less effectively, which we also observed in earlier experiments (Sawyer et al., 2009), suggesting some C-terminal sequences in this fragment are important for actin association.

**Fig 9.**
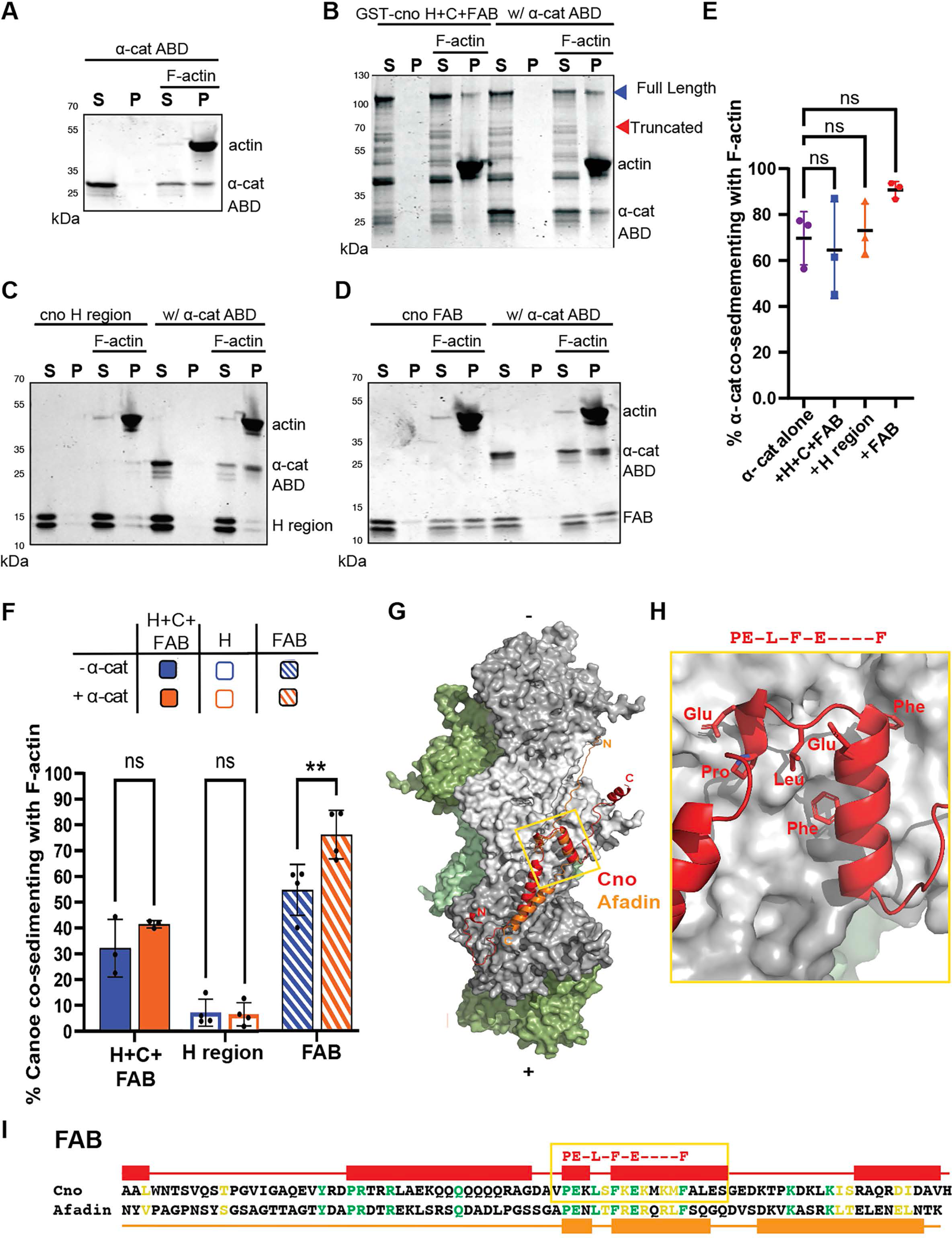
Both the conserved helices and the FAB can bind F-actin. **(A-D)** Immunoblots of the soluble (S) and pellet (P) fractions of different αcat or Cno constructs with or without F-actin. **(A).** The αcat actin-binding domain co-sediments with F-actin. **(B).** A Cno IDR fragment from the conserved helices to the C-terminus (H+C+FAB) co-sediments with F-actin, and this is not enhanced by addition of αcat. **(C).** The conserved helices (H) modestly co-sediment with F-actin, and this is not enhanced by addition of αcat. **(D).** The FAB peptide co-sediments with F-actin, and this interaction is enhanced by addition of αcat. **(E).** Quantification revealing that αcat actin-binding domain co-sedimentation is not substantially enhanced by any Cno peptide. **(F).** Quantification of co-sedimentation of different Cno peptides with or without α-catenin. **(G).** Structural alignment of AlphaFold 3 models of the Cno (red) and Afadin (orange) FAB on actin. One F-actin protofilament is colored in grey, the other in green. **(H).** Zoomed in view of Cno FAB from yellow boxed region in G, showing conserved residues. **(I).** Sequence alignment of the Drosophila Cno and rat Afadin H regions, with identity shown in green font, similarity in yellow font. AlphaFold 3-predicted helices are indicated above or below the respective sequence.

Next, we examined two IDR regions which, in Afadin, were either previously demonstrated to bind F-actin (the conserved helices (H)) or have a name suggesting that binding experiments were done (the FAB). We generated constructs allowing us to express and purify these two peptides (Fig S3, S4A), and first used circular dichroism to examine whether they are helical in solution. As a positive control, we examined Cno’s Dilute domain which produced minima at 208 and 222 nm, well aligned with its predicted secondary structure (Fig S4B; (McParland et al., 2024a). The Afadin H region has a forked helical structure in the F-actin:αcat:Afadin structure (Gong et al., 2025). While many individual helices adopt structure upon binding a partner, we found that the Cno H peptide showed helical character on its own (Fig S4C). To determine if this was due to oligomerization, we analyzed the oligomeric state of the H peptide using SEC-MALS, which revealed a single monomeric species (Fig S4E). In contrast, the FAB peptide had a single minimum at 200 nm, indicative of a disordered peptide Fig S4D). As the FAB has predicted helices, this data may imply the FAB adopts helical character upon binding a partner.

To directly assess the ability to bind F-actin, we tested each fragment for F-actin co-sedimentation. After purification, both the H and the FAB peptides ran as two bands (Fig S5A, C), and we used mass-spectrometry to identify the precise nature of each. The H region peptides were revealed to be a full-length version and one lacking the C-terminal 9 amino acids (Fig S5A, B), while the FAB peptides were full-length and one lacking the C-terminal 28 amino acids (Fig S5C, D). We first focused on quantifying the longer of the two peptides in each case. The H peptide exhibited modest co-sedimentation with F-actin, which was above the background seen in F-actin’s absence, and adding αcat did not enhance H peptide co-sedimentation (Fig 9C, F). In contrast, 55% of the FAB peptide co-sedimented with F-actin and adding αcat significantly increased co-sedimentation (to 76%; Fig 9D, F). While both the full-length and truncated H peptides bound actin similarly (Fig S5E), the longer of the two FAB peptides found F-actin more effectively (Fig S4F), suggesting the 28 C-terminal amino acids removed in the shorter fragment normally enhance actin association.

These results, combined with what is known about Afadin, suggest at least two distinct regions of Cno’s IDR can bind F-actin: the conserved helices (H) and the FAB. ΑlphaFold3 allows us to model protein interactions. We used this tool to model potential interactions of H and the FAB with actin filaments. Cryo-electron microscopy revealed that the H region of Afadin binds both the actin-binding domain of αcat and F-actin, stabilizing this interaction (Gong et al., 2025). In this structure, it forms a forked structure of two α-helices (Fig S6B). The H peptide is reasonably well conserved between mammals and Drosophila and all four residues important for interactions of the mammalian H peptide with αcat and F-actin (Gong et al., 2025) are perfectly conserved (Fig S6A). We used ΑlphaFold3 to model the Cno H peptide bound to a Drosophila actin filament along with the actin-binding domain of Drosophila αcat. This produced a high-confidence model (as assessed by pLDDT; Fig S6D) very similar to that reported for the Afadin H peptide for the region of overlap (Fig S6B vs C). A conserved PW-R motif on the N-terminal helix engaged residues on actin and one αcat molecule, while a conserved LE—F—R motif in the second helix engaged αcat (Fig S6 C,E,F). Thus, we think it is likely that the H peptide interacts with αcat and F-actin in similar modes in both animals.

Next, we used ΑlphaFold3 to model potential interactions of the Drosophila Cno FAB peptide with actin filaments and compared this with a model of the Afadin FAB and F-actin. While the Cno and Afadin predictions varied in the placement of the N- and C-terminal regions of each FAB, both predictions converged with confidence in the placement of a highly conserved FAB region on an actin subunit on the filament (Fig 9G-I; Fig S6G). This FAB helix-turn-helix motif is predicted to use conserved hydrophobic and charged residues to engage features of actin, primarily on subdomain 1 (Fig 9G, H). These data are consistent with our biochemical assays and suggest the FAB indeed is a second actin binding site.

## Discussion

Intrinsically disordered regions (IDRs) play important roles in assembling multiprotein complexes with diverse functions from transcriptional regulation to cell signaling to cell-cell and cell-matrix adhesion. Many proteins in the AJ and tight junction contain IDRs. However, the roles of the IDRs and how their complex sequence structure contributes to their function in vivo remains a key question in the field. Here we address this using the AJ-cytoskeletal linker Cno, defining the roles of its IDR in protein function and localization, and then taking it apart, defining the sequence features important for its biochemical and cell biological functions.

### Cno’s IDR plays important roles in protein function

Multiple junctional proteins include IDRs, but in only a few cases have their functions been explored. The most extensive characterization was of the tight junction scaffold ZO-1. It can phase separate in vitro and in vivo, recruits other tight junction proteins into condensates. The ZO-1 C-terminal IDR is important for tight junction localization and in assembly of functional tight junctions (Beutel et al., 2019). Junctional puncta become more continuous via actin interaction (Beutel et al., 2019) and are retained apically and spread along the apical junction complex by interactions with PATJ (Pombo-Garcia et al., 2024). Likewise, Afadin’s C-terminal IDR can also phase separate in vitro and in vivo and plays a role in localizing Afadin to apical AJs (Kuno et al., 2025). Functional studies Afadin’s IDR (Kuno et al., 2025) and that of Cno (Gurley et al., 2023; Perez-Vale et al., 2021) are more limited, with the studies in cultured MDCK cells suggesting a role in maintaining columnar cell architecture (Kuno et al., 2025).

Drosophila development allowed us to define the mechanistic roles of Cno’s IDR in vivo during morphogenesis. We and others somewhat arbitrarily divided the C-terminal IDR into two regions: the relatively conserved C-terminal region, called the FAB, and the longer, less well conserved region referred to here as the proximal IDR. Our previous analysis revealed that deleting the FAB had no apparent effect on Cno AJ localization, including its enrichment in AJs under tension, suggesting regions outside the FAB may confer F-actin association. Further, the FAB was dispensable for restoring viability and fertility, though sensitized assays revealed it did not retain fully wildtype function (Perez-Vale et al., 2021). Here we deleted the proximal IDR. The effect of this was much more dramatic. *cnoΔProxIDR* mutants were embryonic lethal with strong defects in multiple events of morphogenesis. AJs under tension were destabilized. However, CnoΔProxIDR retained a small amount of residual function, with reduced defects in epidermal integrity relative to the null allele. In this way, the IDR ranked with the RA1 domain as the most critical region of Cno protein we assessed thus far.

### Cno’s IDR is important for AJ localization and its deletion leads to surprising nuclear relocalization

The localization of CnoΔProxIDR was strikingly altered: localization to AJs was substantially reduced, and instead the protein re-localized to nuclei. We initially wondered if this was a biological artifact caused by unforeseen creation of a novel NES. However, recent analysis of Afadin revealed nuclear localization of a construct similar to our CnoΔProxIDR (referred to as AfadinΔC; (Kuno et al., 2025). Although the authors do not mention it, predominant nuclear localization is apparent in Figures 3, 4, and 6. Our data together with these suggest there is a nuclear export signal (NES) in the C-terminal region of the proximal IDR. Two contrasting but not mutually exclusive hypotheses are consistent with this change. First, deleting the IDR may lead to sequestration in nuclei, preventing an otherwise functional protein from localizing to AJs. Second, the AJ may be the preferred site of Cno localization because multivalent interactions with other AJ proteins lead to its retention there. Deleting the proximal IDR may reduce affinity for AJs, releasing Cno and leading to its nuclear import.

We prefer the second hypothesis for several reasons. While CnoIDRΔC localizes prominently to nuclei, it also retains strong localization to AJs and rescues viability and fertility. This suggests nuclear localization per se does not lead to sequestration and thus prevent a protein lacking the NES from localizing to AJs. CnoAfadinIDR also has weak nuclear localization, suggesting its NES may not be fully functional in Drosophila. However, it also rescues viability and fertility, further supporting the idea that some nuclear localization is compatible with Cno function.

There are multiple examples of junctional proteins with known or suspected roles in transcriptional regulation (βcat (Peifer, 1997); p120ctn (Daniel and Reynolds, 1999); ZO-1 (Balda et al., 2003)). Our data raise the question of whether Cno has any normal functions in nuclei. There are predicted nuclear localization signals between the RA domains (Rothenberg et al., 2023) and when we expressed an RA-PDZ protein in Drosophila it localized to nuclei (Bonello et al., 2018). However, we have never observed wildtype Cno in nuclei during any stage of embryogenesis, though we cannot rule out low level accumulation there. Interestingly, there is a long history of reports of Afadin localizing to both AJs and nuclei, beginning with the initial cloning by the Takai lab (Mandai et al., 1997) and continuing in subsequent papers from that lab and others (e.g., (Buchert et al., 2007; Miyahara et al., 2000; Nishioka et al., 2000; Ooshio et al., 2010), but in most cases it was either described as an observation whose significance was not clear or dismissed as an artifact. The short isoform of Afadin lacking the FAB and part of the conserved IDR α-helices is more abundant in nuclei than full length Afadin (Buchert et al., 2007; VanLeeuwen et al., 2014). This aligns with our Cno data revealing that the region C-terminal to the IDR helices is important for nuclear exclusion. There are a few examples in the literature suggesting Afadin has a nuclear role indirectly promoting Histone H3 phosphorylation in response to activating the neurotransmitter receptor NMDA (VanLeeuwen et al., 2014) or the Estrogen receptor (Sellers et al., 2018) and subsequent nuclear translocation of Afadin. Akt-mediated phosphorylation of Serine 1718 in Afadin’s FAB can also trigger nuclear accumulation of Afadin with effects on transcription and cell migration (Elloul et al., 2014; Xu et al., 2015). However, while the Akt recognition site is in a conserved region, Cno has a non-phosphorylatable Alanine in place of S1718. In the future, it will be important to define whether Cno has parallel nuclear roles, perhaps in tissues other than the embryonic ectoderm where most of our work has centered.

### Different regions of Cno’s IDR contribute combinatorially to localization and function

Our previous work revealed the surprising robustness of Cno protein. Proteins individually lacking multiple folded domains or conserved regions, including RA2, the DIL domain, the PDZ, and the FAB, restored viability and fertility, though all were required for full function in sensitized assays (McParland et al., 2024a; McParland et al., 2024b; Perez-Vale et al., 2021). Further, all these mutant proteins localized to AJs, and only the RA domains were essential for mechanosensitive recruitment to AJs under tension. These data supported a model in which Cno is recruited to AJs by diverse multivalent interactions and many individual interactions are not essential for localization or function.

In contrast, CnoΔProxIDR had substantially reduced AJ localization, making the proximal IDR the first Cno region with a clear role in this. To define the underlying mechanism, we took the IDR apart. Our series of smaller IDR deletion mutants reveal that different regions of the IDR act combinatorially to mediate AJ location and Cno function. *cnoIDRΔN, cnoIDRΔH, cnoIDRΔC,* and *cnoΔFAB* are all viable and fertile mutants and all proteins localize effectively to AJs. However, sensitized assays suggest all except *cnoIDRΔH* fail to provide fully wildtype function. Two mutant proteins, CnoIDRΔN and CnoIDRΔC, have partial or strong reduction in enrichment in AJs under tension, consistent with their modest deficits in function. These data further emphasize the resilient and multivalent nature of Cno localization to AJs. In this way, Cno resembles the apical polarity protein Bazooka/Par3, as no Baz protein domain is essential for cortical localization, though each contributes to cortical anchorage in a specific manner (McKinley et al., 2012).

Given this, what are the precise roles of each region of the IDR—N, Helices, C, and FAB— and how do they contribute to function? The conserved helices have well defined biochemical properties in Afadin—cryo-EM reveals they bind F-actin and the actin-binding domain of αcat to stabilize this interaction. Sequence conservation, F-actin binding assays, and modeling are consistent with these helices playing a similar role in Drosophila Cno. Thus, we were quite surprised at the modest deficits in function seen in *cnoIDRΔH,* which was the least affected of our mutants. Further, the *cnoIDRHelixOnly* mutant reveals the conserved helices are not sufficient for function, while its slight “dominant-negative” phenotype suggests the helices alone can disrupt function of AJs, perhaps allowing CnoIDRHelixOnly to bind some, but not all, of its necessary protein partners thus sequestering them and further disrupting AJ-cytoskeletal linkage. The FAB, despite its name, had not been previously demonstrated to bind F-actin. Our data reveal it does so, and that its interaction with F-actin is further enhanced by αcat. However, like *cnoIDRΔH*, *cnoΔFAB* is also viable and fertile. One possibility is the conserved helices and the FAB act in a redundant fashion, with each interacting with F-actin and perhaps αcat. In the future it will be important to test a mutant removing both regions.

The functional roles of the IDR N- and C-terminal “spacers” are also intriguing. Each spacer differs in length, sequence composition and charge between mammalian Afadin and Cno. The N-terminal spacer has no sequence identity between Afadin and Cno, while the C-terminal spacer only shares a single short, conserved motif. One role they could play is simply to provide space/volume between more conserved “stickers” like the conserved helices and FAB. Consistent with this, the Afadin IDR largely replaced Cno function, at least to the extent of restoring viability and fertility. Neither spacer is individually required for restoring viability or fertility, and thus their individual “spacer” functions are not essential. However, deleting both led to a strong reduction in Cno function, consistent with roles as spacers and/or functional redundancy in interactions with binding partners. In the future, it would be interesting to scramble the sequence of one or both spacers and see if function is retained—this would help distinguish between roles as spacers or embedded sequence motifs.

Our work also revealed two clear functions of the “spacers”. First, the C-terminal spacer appears to contain an NES, as deleting this region led to nuclear accumulation. Despite the limited sequence identity in this region, the Afadin IDR largely but not completely restored nuclear exclusion. Second, and more surprising, the C-terminal region is critical for Cno’s mechanosensing properties. Deleting it, or replacing it with that region from Afadin, virtually eliminated enrichment of mutant Cno proteins at TCJs. This loss correlated with reduced function in our sensitized assays, but, to our surprise, proteins lacking this mechanosenstive recruitment could still restore viability. The only other mutants altering altered TCJ enrichment were mutations in the RA-domains (McParland et al., 2024b; Perez-Vale et al., 2021)—future work to define the mechanisms of mechanosensing will be important.

## Acknowledgements

We are grateful to the Drosophila Genomics Resource Center (NIH Grant 2P40OD010949) for the Drosophila alpha-catenin clone, Dr. Ashutosh Tripathy of the UNC Macromolecular Interactions Facility, Dr. Nat Prunet of the Biology Imaging Core, Rachel Szymanski for help getting this project off the ground, and to Peifer, Bergstralh, Finegan, and Williams lab members for helpful discussions. This work was funded by NIH R35 GM118096 to M. Peifer.

## Methods

### Cloning of cnoΔIDR constructs

Most constructs were generated using Azenta Life Science (Waltham, MA, USA) cloning and mutagenesis service. Details and sequences arein Fig S1. *cnoΔIProxDR* was generated by removing all nucleotides corresponding to amnio acids M1130-Q1926 in the Canoe protein using *cnoWT-GFP* as the starting construct (Perez-Vale et al, 2021). Then using Azenta Life Science’s gene synthesis service, we generated a small sequence containing a 5’ SbfI RE site and a 3’ SpeI RE site with a small Gly-Gly linker to maintain the proper reading frame (5’-GCCTGCAGGGGCACTAGT -3’) and subcloned that into the deletion generated above.

This new cnoΔProxIDR-SbfI-SpeI construct was then used to generate all the subsequent mutants via RE cloning using those SbfI and SpeI overhangs. This construct was verified by PCR (ΔProxIDR forward primer 5’ – GCGTGGTAATCGCCGTATGTC – 3’ and ΔProxIDR reverse primer 5’ – CAACACAATCTCGACGCCT – 3’) to confirm sequence and further verified by RE digest and western blot analysis. *cnoIDRΔN* was generated in a similar manner. deleting all nucleotides corresponding to amino acids M1130-Y1566, *cnoIDRΔC* removed to amino acids I1673-Q1926, *cnoIDRΔH* removed to amino acids V1567-A1672, and *cnoIDRHelixOnly* removed all to amino acids except those between V1567-A1672. All of these constructs were generated. The *cnoCanoeIDR* construct was generated by adding back amino acids M1130-Q1926 into the SbfI-SpeI site resulting in a final product that contained linker amino acids PAG upstream and GTS downstream of the endogenous IDR sequence. *cnoAfadinIDR* was generated similarly by adding nucleotides corresponding to amino acids A1097-A1750 of the rat Afadin protein into the SbfI-SpeI sites. All constructs were verified by DNA sequencing. Each construct also contains the *w^+^* selectable marker and is flanked by attR and attL sites allowing site-specific integration into the attP site at the cnoΔΔ locus as described in Perez-Vale et al., 2021. These constructs were then sent to BestGene (Chino Hills, CA, USA) for embryo injections. Briefly, DNA was injected into *PhiC31/int^DM. Vas^; cno*ΔΔ embryos (BDSC stock 94023). F1 offspring were screened for the presence of the *w^+^* marker and outcrossed to *w; TM6B, Tb/TM3, Sb* (BDSC stock 2537) to generate a balanced stock over TM3. Each stock was verified by PCR, sequencing and western blot analysis. We outcrossed these stocks to a *yw* stock with a wild type 3^rd^ chromosome to remove potential passenger mutations from the mutant chromosomes. This occurred for multiple generations selecting for the linked *w^+^* marker in each generation. We were thus able to generate homozygous stocks for some of our mutant alleles.

### Fly Work

All experiments were performed at 25°C unless noted otherwise. Flies of the *yellow white* genotype [Bloomington Drosophila Stock Center (BDSC), stock 1495] were used as controls and are referred to in the text as wildtype (WT). Maternal/zygotic mutants of *cnoΔProxIDR* and *cnoIDRHelixOnly* were made using the FRT/ovoD approach. Briefly, third-instar larvae generated by crossing *cno*ΔProx*IDR/TM3* or *cnoIDRHelixOnly/TM3* virgin females to *P{ry[+t7.2]=hsFLP}1, y[1] w[1118]; P{neoFRT}82B P{ovo−D1−18}3R/TM3, ry[*], Sb[1]* males were heat-shocked in a 37°C water bath for 2 hours each on two consecutive days. Then, virgin female progeny with the germline genotype *hsFLP1; P{neoFRT}82B P{ovo−D1−18}3R/P {neoFRT}82B cnoΔProxIDR* were collected and subsequently crossed with *cnoΔProxIDR/TM3 Sb* males. The embryos generated from this cross were analyzed. Maternal/zygotic mutants identified by absence of staining with our Cno antibody which recognizes an epitope in the proximal IDR, and referred to in the text as simply *cnoΔProxIDR*. Sensitized assays were done using mutant flies that are heterozygous for each *cno-mutant* allele with the null allele *cno^R2^* as described in Perez-Vale et al, 2021.

### Immunoblotting and quantification

Table S3 lists the antibodies and dilutions used for these experiments. We determined accumulation levels wildtype Cno, GFP-tagged mutants, and α-tubulin by immunoblotting embryonic lysates that were collected at 4-8 hours after egg laying. Embryonic lysates were generated as in (McParland et al., 2024b). Briefly, embryos were collected into 0.1% Triton X-100, dechorionated for 5 minutes in 50% bleach, and washed three times with 0.1% Triton X-100. Next, lysis buffer (1% NP-40, 0.5% Na deoxycholate, 0.1% SDS, 50 mM Tris pH 8.0, 300 mM NaCl, 1.0 mM DTT, HaltTM Protease and Phosphatase Inhibitor Cocktail (Thermo Fisher Scientific, #78442) (100×), and 1 mM EDTA) was added, embryos were ground with a pestle for ∼20 seconds and then placed on ice. After 10 minutes on ice, embryos were once again ground with a pestle for ∼20 seconds and subsequently centrifugated at 16,361xg for 15 minutes at 4°C. Protein concentration was assessed using the Bio-Rad Protein Assay Dye, recording absorbance at 595 nm with a spectrophotometer. Protein lysates were run on 7% SDS-PAGE, and proteins were transferred onto nitrocellulose membranes.

Membranes were blocked using 10% bovine serum albumin (BSA) diluted in Tris-buffered saline with 0.1% Tween-20 (TBST) for 1 hour at room temperature. Primary and secondary antibodies were diluted in 5% BSA diluted in TBST. Primary antibody incubations were performed overnight at 4°C, and secondary antibody incubation was performed for 45 minutes at room temperature. We used the Odyssey CLx infrared system (LI-COR Biosciences) to image the membranes, and band densitometry was carried out using Empiria Studio® Software (LI-COR Biosciences).

### Embryo fixation and immunofluorescence

Embryo fixation and immunofluorescence were performed as previously described (Noah or Emily papers). Briefly, flies were placed into cups containing apple juice agar plates with added yeast paste. Embryos were then collected using a paint brush and 0.1% Triton-X-100. They were then bleached to remove the chorion layer using 50% bleach and washed three times with 0.03% Triton X-100 with 68 mM NaCl. They were then fixed in 95°C in a 0. 03% Triton X-100 with 68 mM NaCl and 8 mM EGTA salt solution for 10 seconds, and brought back down to temp on ice for 30 minutes. The vitelline membrane was removed by vigorous shaking in a 1:1 solution of n-heptane and 95% methanol:5% EGTA and then washed three times with a 95% methanol and 5% EGTA solution. To prepare for staining, embryos were washed three times with 5% normal goat serum (NGS; Thermo Fisher Scientific) and 0.1% saponin in phosphate-buffered saline (PBS) (PBSS-NGS) and then blocked for an hour using 1% NGS in PBSS-NGS. Antibodies were diluted (see antibody dilution table below) in a PBS solution containing 1% bovine serum albumin and 0.1% saponin and incubated overnight at 4°C. Embryos then received three 15-minute washes with PBSS-NGS and incubated in secondary antibodies for 2 hours at room temperature. Embryos were then mounted on glass slides using a homemade Gelvatol solution (recipe from the University of Pittsburg’s Center for Biological Imaging).

### Image acquisition and analysis

Embryos were imaged using a 40X/NA 1.3 Plan-Apochromat oil objective on a LSM 880 confocal laser-scanning microscope (Carl Zeiss, Jena Germany) running on ZEN Black 2009 software. Using Photoshop (Adobe, San Jose, CA) all images were adjusted to have uniform input levels, brightness, and contrast.

### Automated TCJ/multicellular junction enrichment analysis

We performed membrane segmentation using the EpySeg graphical user interface (GUI) (Aigouy et al., 2020), which employs a pre-trained deep learning model for image segmentation. Specifically, we utilized the "v2" model from the segmentation_models library (https://github.com/qubvel/segmentation_models), with the following architectural specifications: Model Architecture: Linknet, Backbone Network: VGG16, Activation Function: Sigmoid, and Input Image Dimensions: Dynamically adjusted (0 width and height). The segmentation was performed on representative images, with predictions generated based on the Armadillo staining channel of de-identified images to achieve a single-blinded analysis. After membrane segmentation, we used FIJI (ImageJ) and the Analyze Skeleton plugin (Arganda-Carreras et al., 2010) (https://github.com/fiji/AnalyzeSkeleton) to identify the intersections of tricellular and multicellular borders. The intersection points were then isolated and used as reference points of interest for subsequent analysis. We next used the ModularImageAnalysis (MIA) software developed in the Wolfson Bioimaging Facility at the University of Bristol (Cross et al., 2024). This software allows for fully unique and customizable workflows and analysis. Using this, we loaded the segmented Armadillo image from EpySeg as the skeleton marking all cell borders. We also inputted the location of tri- and multi-cell borders from FIJI. We then preformed two iterations of dilations of these points to make a larger area of the border intersections. This larger area was then subtracted from the skeleton to give us disconnected bi-cells with the ends of the bi-cell edges no longer overlapping the cell-intersection area determined previously and with sufficient separation of signal for larger tri-cell junctions. Lastly, we used the GFP channel as the input. We preformed manual thresholding of the signal, elimination of background signal/noise, and manually selected a 512 pixel by 512 pixel area to perform the analysis on (representing one-quarter of the full 1024 image). Using the GFP signal we measured the signal intensity along each bi-cell border as well as its associated tri- and multi-cell junction and compared the ratio of these two intensity values to determine the enrichment at tri-cellular junctions.

### Analysis of junctional gaps

For each genotype, we selected closeups of similar magnifications of seven stage 7 and seven stage 8 embryos stained to visualize Arm, which outlines AJs, and selecting lateral views from the ventral midline toward the amnioserosa. These were then coded, randomized, and scored blind for gaps in tricellular or multicellular junctions, or places where the junctional protein signal was broadened. Samples were then unblinded, and total gap number was then recorded per field of view.

### Statistics and quantification

Graphs were made using GraphPad Prism version 10.4.1 (GraphPad Software, Boston, MA, USA). For box and whisker plots, boxes represent 25th–75th percentile ratios and whiskers represent the 5th–95th percentiles. Statistical analyses were performed in Prism using Welch’s unpaired *t*-test or Brown–Forsythe and Welch ANOVA test and unless otherwise noted, error bars represent 95% Confidence Intervals. Kolmogorov-Smirnov analysis was used for the comparison of GFP levels between CnoIDRΔC and CnoIDRAfadin compared to CnoWT-GFP.

### Cuticle preparation and analysis

Embryonic cuticles were prepared according to (Wieschaus and Nüsslein-Volhard, 1986). Females were placed in cups and eggs were collected on apple juice agar plates with yeast for less than 24 hours at 25°C. Eggs were then aligned on a fresh apple juice agar plate without yeast and incubated at 25°C for 48 hours to allow embryos to develop fully and viable embryos to hatch. Unhatched embryos were collected in 50% bleach and dechorionated for 5 minutes. The bleach was removed and they were transferred to glass slides into 100µl of 1:1 Hoyer’s medium:lactic acid. The slides were incubated at 60°C for 24-48 hours. They were then stored at room temperature. Images were taken using a Nikon Labophot with a 10x Phase 2 lens, and captured on an iPhone, and placed into categories based on morphological criteria, including counting unfertilized eggs and removing them from the total. At least two separate crosses were used for each genotype.

### Scoring eye phenotypes

Flies were maintained at 25℃ on nutrient-rich Drosophila media and prepupae selected and stored in humidified chambers until dissection at 40h after puparium formation (APF). At 40h, pupae were submerged in a droplet of 1.5x PBS, removed from the pupal casing, and decapitated. The head casing was then removed to expose the brain complex, and it was rinsed in 1.5x PBS twice to remove any excess fat and tissue. The brain complex was then fixed in 3.7% formaldehyde for 20 min, washed in 1.5x PBS twice, and blocked in 1.5x PBST for 30 min. The tissue was then placed in rat anti E-Cadherin (1:50; DSHB) and rat anti N-Cadherin (1:50; DSHB) overnight at 4℃ before being rinsed in 1.5x PBS twice and blocked in 1.5x PBST for 30 min. The tissue was then stained with Goat anti-Rat 568 secondary (1:100; Life Tech) to detect AJs and retinas and imaged with a Zeiss LSM 980 with Airyscan 2 using a 63x/1.4 oil Plan Apochromat objective. Patterning errors were scored in 10 eye discs per genotype spanning 77-115 ommatidia per genotype. Data were analyzed for statistical significance using PRISM, normality was determined using the Kologorov-Smirnov test, and significance was calculated using an ordinary one-way parametric ANOVA. Images were processed for publication using FIJI and Adobe Photoshop.

### Cloning and purification of Canoe fragments and the Alpha Catenin actin binding domain

DNA encoding the *Drosophila melanogaster* Cno H region (residues 1567-1670) and Cno FAB (residues 1967-2051) were generated using PCR (H region forward primer, 5’-GGCAGGACCCATATGGCGTCCAATCAAGGGAACAATCGTC; H region reverse primer, 5’-GCCGAGCCTGAATTCTTAactgacctgaccgccgccagccaatc-3’; FAB forward primer, 5’-GACTATCGTCATATGGCTGCCCTCTGGAACAC-3’; FAB reverse primer, 5’-GACTATCGTCTCGAGTTAGTGCACCGCCGCGTCTATATC-3’;) and sub-cloned into pET28 (Millipore Sigma, Burlington, MA) using Nde1 and EcoR1 restriction enzymes and T4 DNA ligase (New England Biolabs, Ipswich, MA).

DNA encoding *Drosophila melanogaster* Cno H+C+FAB (residues 1560-2051) was generated using PCR (H+C+FAB forward primer, 5’-CGCTGTTCGGGTACCATGGGATCCCCGCCAAAGGGCAGCTATGTGGCGTCC-3’; H+C+FAB primary reverse primer, 5’-GCCGAGCCTGAATTCTTAGTGCACCGCGTCTATATCTCGTTG -3’; secondary reverse primer to append a C-terminal His_6_ tag, 5’-GCCGAGCCTGAATTCTTAGTGGTGATGATGGTGATGGTGCACCGCGTCTATATCTCG-3’) and sub-cloned into pGEX-6p2 using BamH1 and EcoR1 restriction enzymes and T4 DNA ligase (New England Biolabs).

Full length *Drosophila* alpha-catenin cDNA was obtained from the DGRC on a Whatman FTA disc, stock number FI19832 (DGRC Stock 1649238; https://dgrc.bio.indiana.edu//stock/1649238 ; RRID:DGRC_1649238) and was used as the template for PCRs that amplified the *Drosophila* alpha-catenin long ABD (*Dm* long α-cat: residues 667-917) (*Dm* long α-cat forward primer, 5’-CGCTGTTCGGCTAGCGATCAAACCGTTGACGAATATCCCGATATAAG-3’; Reverse primer 5’-CACTCGTTCAAGCTTTTAAACAGCGTCAGCAGGACTCTGG-3’) coding region, which was then sub-cloned into pET28 (Millipore Sigma, Burlington, MA) using NheI and HindIII restriction enzymes and T4 DNA ligase (New England Biolabs, Ipswich, MA). All plasmids were sequence verified.

The plasmids were transformed into *Escherichia coli* BL21 DE3 pLysS cells and grown to an optical density at 600 nm of 0.8 in medium containing antibiotic (50 µg/l kanamycin or 100 µg/l ampicillin for respective vectors) at 37°C. The temperature was lowered to 20°C (excluding the GST-Cno H+C+FAB-H_6_ construct, which was kept at 37°C), and protein expression was induced with 100 µM IPTG for 16 h. Cells were harvested by centrifugation, resuspended in buffer A [25 mM Tris-HCl pH 8.5, 300 mM NaCl, 10 mM imidazole and 0.1% β-mercaptoethanol (β-ME)] at 4°C and lysed by sonication (the GST-Cno H+C+FAB buffer A was supplemented with 1% Triton X-100 and 10% glycerol). Phenylmethylsulfonyl fluoride was added to 1 mM final concentration. Cells debris was pelleted by centrifugation at 17,000 x g for 1 h. Supernatants for constructs cloned in the pET28 vector were loaded onto a 10 ml Ni^2+^-NTA column (Qiagen, Hilden, Germany). The column was washed with 1 L buffer A, and the protein batch eluted with 100 mL buffer B (buffer A supplemented with 290 mM imidazole).

Supernatant for the GST-Cno H+C+FAB-H_6_ construct was loaded onto a 10 ml Glutathione Sepharose 4 Fast Flow column (Cytiva, Marlborough, MA). The column was washed with 1 L of buffer N [25 mM Tris-HCl pH 8.5, 300 mM NaCl, and 0.1% β-ME] and the protein batch eluted with 100 mL buffer N supplemented with 50 mM Glutathione. Eluate was then loaded onto a 10 ml Ni^2+^-NTA column, washed, and eluted as described above.

N-terminally His-tagged proteins received 5 µl (1.2 mg/ml) of bovine α-thrombin (Prolytix, Essex Junction, VT) and 1 mM CaCl_2_, and were incubated at room temperature for 3 hours and then 4°C for 48 h to proteolytically cleave off the N-terminal His_6_ tag. Protein was then filtered consecutively over 0.5 ml benzamadine sepharose (Cytiva, Marlborough, MA) and 10 ml Ni^2+^-NTA resin (Qiagen, Hilden, Germany), and the flowthrough was collected.

All constructs were dialyzed against 4L buffer N for 20 hours using 3.5K molecular weight cutoff (MWCO) SnakeSkin dialysis tubing (Thermo Fisher Scientific, Waltham, MA). Protein was concentrated in a Millipore Sigma 3K MWCO centrifugal concentrator, aliquoted and stored at −80°C. Canoe Dilute Domain (aa 613-1006) cloning and purification was described in McParland, et al. 2024

### F-actin co-sedimentation assays

F-actin co-sedimentation assays were adapted from Gong et al., 2025 (as follows: Lyophilized rabbit skeletal muscle actin (Cytoskeleton, Inc., Denver, CO) was reconstituted to 10 mg/ml (233 μM) with 100 μL deionized water and then diluted to 20 μM in Actin Buffer (2 mM Tris pH 8.0, 0.1 mM CaCl_2_, 0.2 mM ATP, 0.5 mM DTT) and stored at -80°C in 112 μL aliquots. Actin was polymerized by mixing 112 μL of thawed monomeric actin with 12.6 μL 10x KMEI buffer (10X = 500 mM KCl, 10 mM MgCl_2_, 10 mM EGTA, 100 mM imidazole pH 7.0) supplemented with 5 mM ATP and incubated at room temperature for 1 hour (final Actin concentration of 18 μM). Canoe fragment constructs were diluted in 1X KMEI buffer to a final concentration of 13.5 μM, either alone or mixed with 9 μM alpha-catenin ABD construct, and pre-cleared by ultracentrifugation at 80,000 rpm (278,000 x g) in a TLA-100 rotor (Beckman Coulter, Brea, CA) for 15 minutes at 4°C. 31.1 μL of the supernatant was mixed with either 38.9 μL of 18 μM polymerized actin or 38.9 μL of 1X KMEI buffer to give final concentrations of Canoe at 6 μM, alpha catenin at 4 μM, and actin at 10 μM. Mixtures were incubated for 30 minutes at room temperature, then ultracentrifuged at 80,000 rpm (278,000 x g) for 30 minutes at 4°C. ¾ of the supernatant (52.5 μL) was mixed with 17.5 μL of 4x SDS-PAGE loading buffer. The remaining ¼ of the supernatant was removed, the pellet was washed twice with 1X KMEI buffer, and then dissolved in 4/3 volume of 1x SDS-PAGE loading buffer (93.3 μL). Samples were separated by SDS-PAGE, stained with Coomassie Brilliant Blue, scanned on a LI-COR Odyssey scanner (LI-COR Biosciences, Lincoln, NE), and quantified with FIJI.

### Size exclusion chromatography - multi-angle light scattering

A 100-μl sample of the *D.m.* Canoe H region construct (aa 1567-1670; 231 μM) was injected onto a Superdex 200 10/300 GL gel filtration column (Cytiva, Marlborough, MA) in 25 mM Tris pH 8.5, 100 mM NaCl, 0.1% β-ME, 0.2 g/L sodium azide, and run in-line with a Wyatt DAWN HELIOS II light scattering instrument and a Wyatt Optilab t-rEX refractometer (Waters | Wyatt Technology, Santa Barbara, CA). Molecular weight was calculated with light scattering and refractive index data using the Wyatt Astra V software package (Waters | Wyatt Technology Corp.). SECMALS data presented are representative of experiments conducted in duplicate.

### Circular dichroism

*D.m.* Canoe constructs (Dilute domain (aa 613-1006); IDR H region (aa 1567-1670); FAB (aa 1967-2051)) were exchanged into 10 mM sodium phosphate, pH 7.4, 50 mM sodium fluoride and diluted to a final concentration of 0.2 mg/ml in 300 μl volume. Initial circular dichroism spectra of constructs were collected at 20°C using a Chirascan-V100 spectrometer (Applied Photophysics, Leatherhead, UK). Spectra were recorded from 260 to 185 nm with a step size of 0.5 nm using a 1 mm-path-length cuvette. The time per point was maintained at 1.25 sec. CD melt spectra were obtained, recording values at 208 and 220 nm (as well as 200 nm for the FAB construct) in 1°C steps from 20-94°C with the time per point maintained at 1.25 sec. After each melt, a final spectrum was recorded from 260 to 185 nm at 94°C for each construct. A base-line CD spectrum of the buffer alone was taken and subtracted from each spectrum. CD data presented are representative of experiments conducted in duplicate.

### AlphaFold 3 structural modeling

Structural modeling was performed using AlphaFold 3 via alphafoldserver.com (Cross et al., 2024). To generate a model of the Cno H region bound to F-actin in the presence of the high-affinity alpha-catenin actin binding domain region, the following were input to AlphaFold 3: *D.m.* actin 5C (6 copies), 6 ATP molecules, 6 Mg^2+^ ions, alpha-catenin (aa 719-917, 3 copies), *D.m.* Canoe (aa 1567-1670, 2 copies). The best model produced an F-actin structure composed of two three-subunit protofilaments with three alpha-catenin actin binding domains and two Canoe H regions bound, similar to the 9dva cryo-EM structure of the mammalian homologs (Gong et al., 2025). The composition of the model presented in figures was trimmed to show five actin subunits (bound to ATP and Mg^2+^), two alpha-catenin actin binding domains, and one Canoe H region, akin to the chains presented in the 9dva structure. To generate a model of the Cno FAB region bound to F-actin, the following were input to AlphaFold 3: *D.m.* actin 5C (6 copies), 6 ATP molecules, 6 Mg^2+^ ions, *D.m.* Canoe (aa 1934-2051, 1 copy). The best model produced an F-actin structure composed of two three-subunit protofilaments with one Canoe FAB region bound. To generate a model of the Afadin FAB region bound to F-actin, the following were input to AlphaFold 3: *Rattus norvegicus (R.n.)* beta-actin (6 copies), 6 ATP molecules, 6 Mg^2+^ ions, *R.n.* Afadin (aa 1712-1829, 1 copy). The best model produced an F-actin structure composed of two three-subunit protofilaments with one Afadin FAB region bound. PyMOL (Schrödinger, New York, NY) was used for structural alignment and image generation.

## SUPPLEMENTAL FIGURES

**Fig S1.**
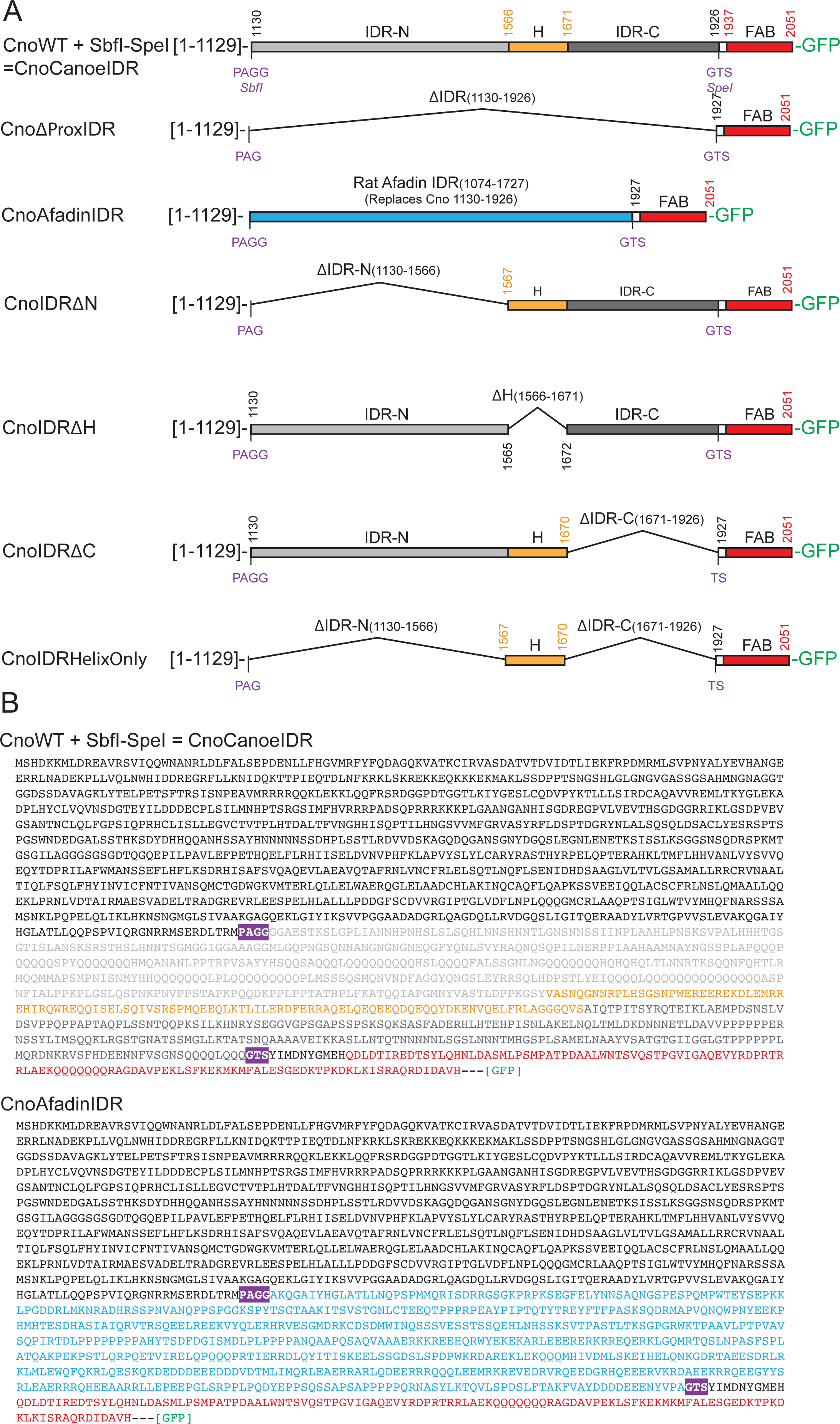
Design of *cno* rescue constructs. **(A)**. Proteins encoded by the *cno* structure/function mutant constructs used in the current study. Amino acids introduced due to the insertion of *SbfI* and *SpeI* restriction enzyme sites and subsequent cloning strategies are indicated in purple font. Cno regions are color-coded as follows: IDR-N (light grey), H region (orange), IDR-C (dark grey), FAB (red). The rat afadin IDR is colored blue. All constructs have a C-terminal GFP. **(B).** The full sequences of CnoCanoeIDR and CnoAfadinIDR.

**Fig S2.**
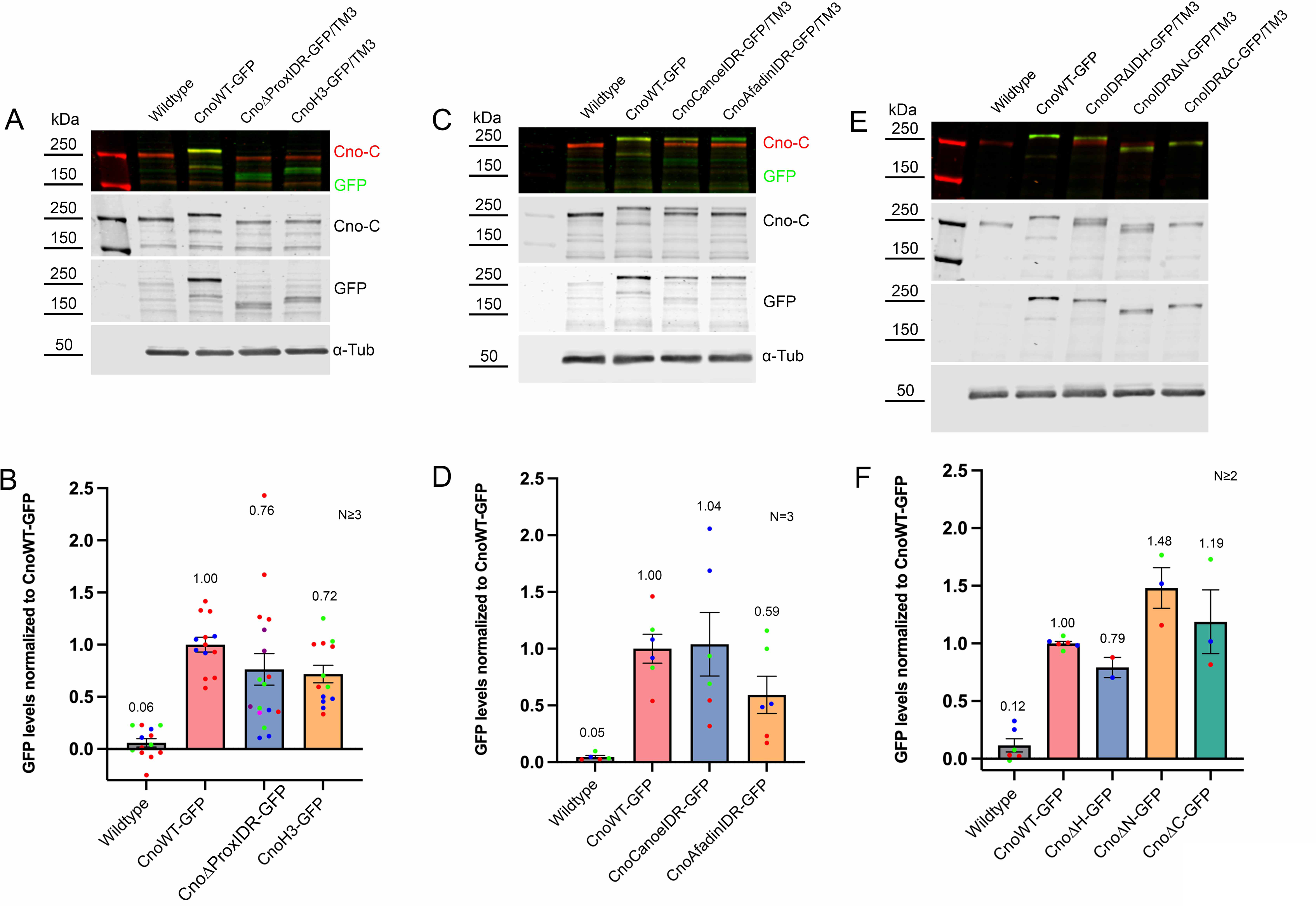
Cno mutant proteins accumulate at roughly wildtype levels. **(A, C, E).** Representative images of immunoblots from embryos of the indicated genotypes, using protein extracts from 4-8 hour old embryos. Following SDS-PAGE, proteins were transferred onto nitrocellulose membranes and immunoblotted with antibodies against the C-terminal region of Cno, GFP, and α-tubulin (loading control). *y w* (our wildtype control) and homozygous *cnoWT-GFP* embryos served as negative and positive controls for the GFP antibody, respectively. The other mutant samples are from parents heterozygous for the mutant allele and a Balancer chromosome wildtype for *cno*. Protein ladders are shown in the left-most well of each panel, and the corresponding molecular weights are indicated. **(B,D, F).** Quantification of expression levels of Cno mutant proteins relative to CnoWT-GFP, the levels of which are normalized to 1. Since mutant protein extracts were generated from heterozygous animals, expression values for mutant protein levels were multiplied by two as a proxy for homozygous expression, and these values are represented in the graphs. Dots of different colors indicate separate biological replicates whereas dots of the same color within each bar represent technical replicates (i.e., single gel with 2 wells containing the same biological samples or different gel using same biological sample). N=number of mutant biological replicates. Top line of column depicts the mean value, and narrower bands indicate standard error of the mean.

**Fig S3.**
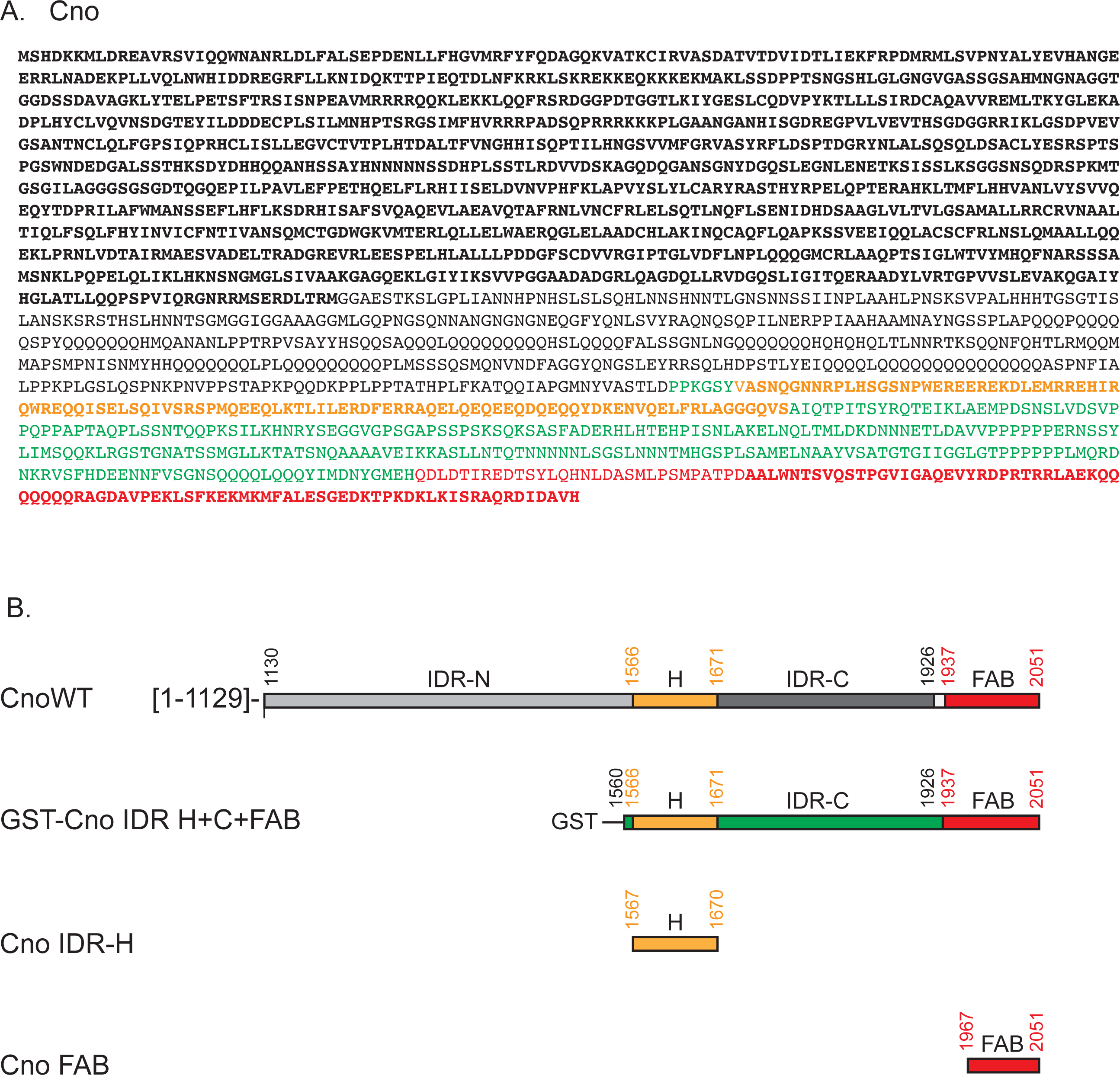
Cno bacterial expression constructs. **(A).** Full length *Drosophila* Canoe sequence. Cno IDR regions are delineated in the image at bottom, as shown in Fig S1**. (B).** Cno bacterial expression constructs. GST-Cno H+C+FAB_1560-2051_-H_6_ (in pGEX-6p2) is color-coded as in A. The purified construct retains the GST and H_6_ affinity tags. Cno H Region_1567-1670_ and Cno FAB_1967-2051_ (in pET28) correspond to the bold orange and red sequence in A. The purified Cno H and FAB constructs had their N-terminal H_6_ tags proteolytically removed.

**Fig S4.**
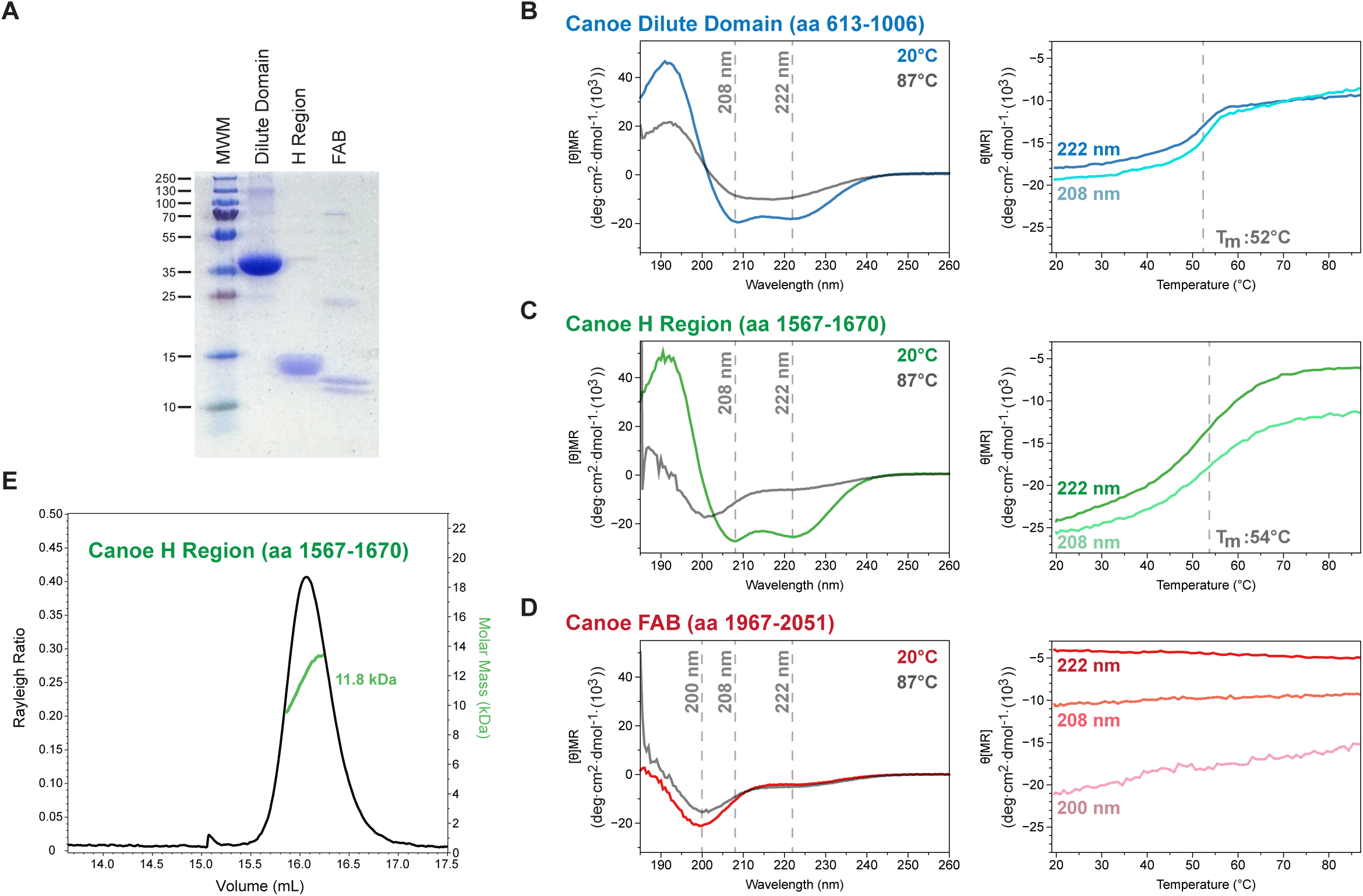
The Cno H region is monomeric and helical while the FAB is disordered. **(A).** SDS-PAGE of purified Cno Dilute domain, H region, and FAB. **(B, C).** Circular dichroism (CD) spectra (left) of the Cno Dilute domain (B), and the H region (C) display minima at 208 and 222 nm, indicative of α-helical structure. Thermal melts (right) of these constructs, monitored at 208 and 222 nm, reveal temperature-dependent structure, with T_m_ values of 52°C (Dilute domain) and 54°C (H region). **(D).** CD spectra (left) and thermal melt (right) of Cno FAB reveals no ordered secondary structure. **(E).** SEC-MALS analysis of the Cno H region results in a single dominant peak with an experimental molecular mass of 11.8 kDa. The formula mass is 12.9kDa, and mass spec identified a second, C-terminal truncated species at 12.1 kDa (Fig S5A), indicating that the Cno H region is monomeric in solution.

**Fig S5.**
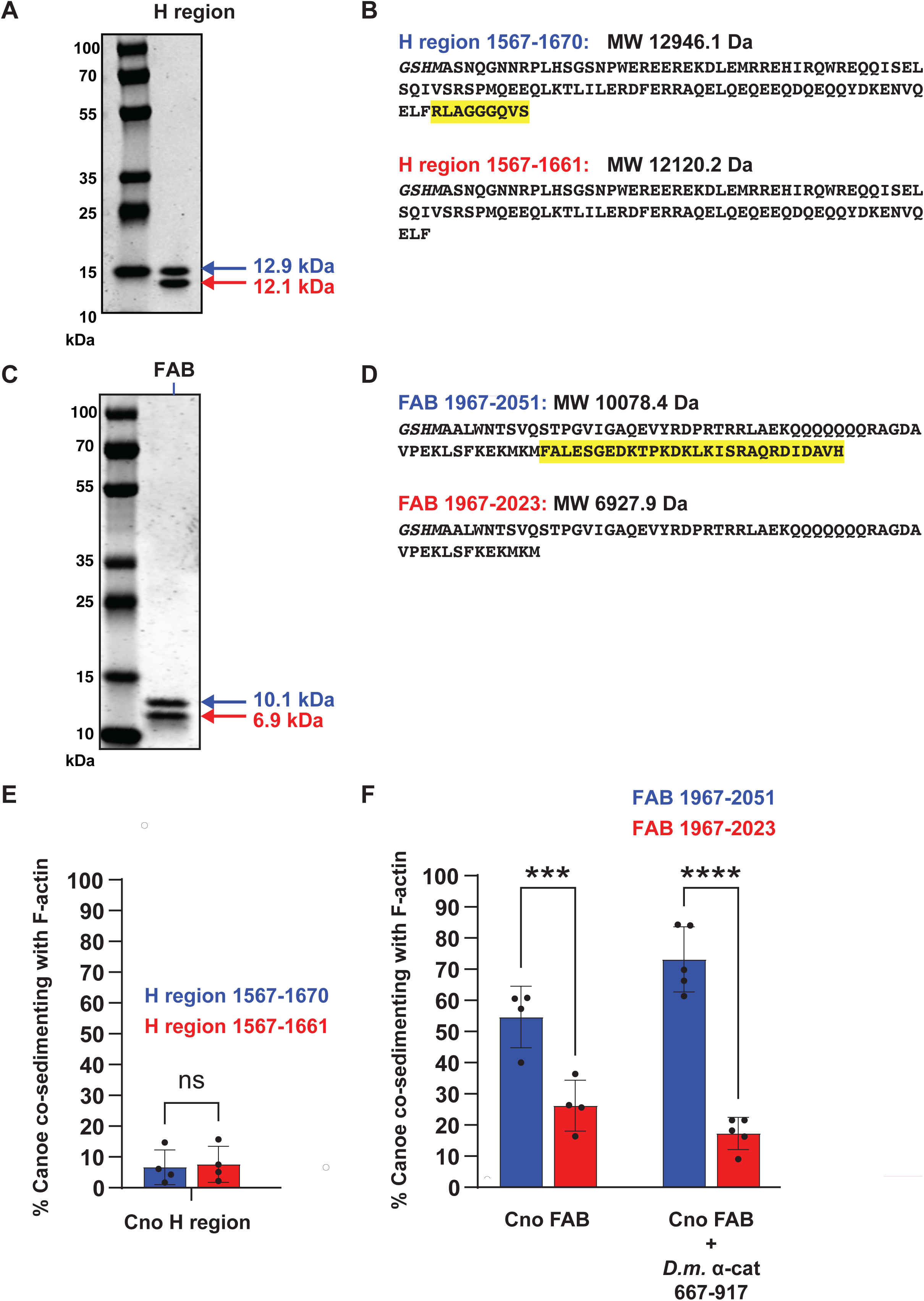
Mass spectrometry reveals C-terminal truncations of the purified Cno H region and the FAB. **(A)**. SDS-PAGE gel of the purified Cno H region yields a doublet. Mass spectrometry analysis identified two major species at 12.9 and 12.1 kDa, corresponding to the predicted mass of a C-terminal 9-residue truncation. **(B).** Sequences for the full-length Cno H region construct (aa 1567-1670) and the predicted sequence of the truncated species (aa 1567-1661) are indicated along with their formula weights. **(C).** SDS-PAGE gel of the purified Cno FAB yields a doublet. Mass spectrometry analysis identified two major species at 10.1 and 6.9 kDa, corresponding to the predicted mass of a C-terminal 18-residue truncation. **(D).** Sequences for the full-length Cno FAB construct (aa 1967-1670) and the predicted sequence of the truncated species (aa 1567-1661) are indicated along with their formula weights. **(E).** Plot showing the relative percentage of each purified Cno H region species co-sedimenting with F-actin. **(F).** Plot showing the relative percentage of each purified Cno FAB species co-sedimenting with F-actin alone, or in the presence of *Drosophila* alpha-catenin.

**Fig S6.**
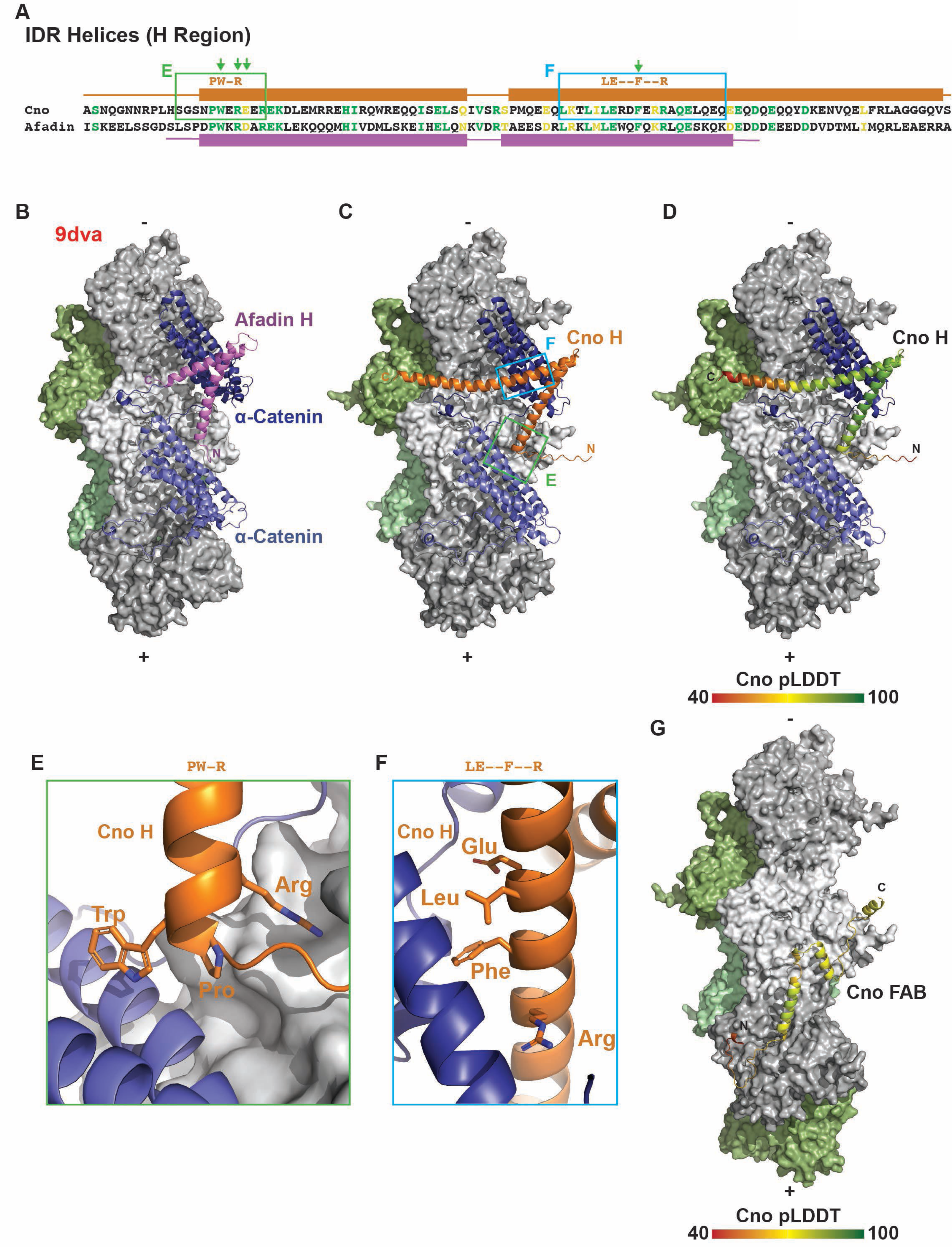
The Cno H region and FAB are predicted to engage F-actin. (**A).** Sequence alignment of the *Drosophila* Cno and mouse Afadin H regions, with sequence identity shown in the green font, and similar amino acids in the yellow font. AlphaFold 3-predicted helices and modeled helices from the 9dva structure are indicated above and below the respective sequence. **(B).** Cryo-EM structure of the mammalian Afadin – αcat– F-actin structure (pdb accession code 9dva). The Afadin H region is shown in purple. One F-actin protofilament is colored in grey, the other in green. **(C).** AlphaFold 3-predicted structure of the *Drosophila* Cno H region – αcat– F-actin complex reveals a similar architecture as the mammalian homologs shown in B. The Cno H region is shown in orange. **(D).** Predicted structure as presented in C, with the Cno H region color-coded according to pLDDT confidence scale presented. **(E-F)**. Zoomed view of Cno H region conserved residues interacting with αcat and F-actin (sequence regions boxed in A). **(G).** Predicted structure as presented in Fig 9G, shown with the Cno FAB color-coded according to the pLDDT value confidence scale.

| Genotype | VF OK | Mild defects | Moderate defects | Wide open | Total | Percent defective |
| --- | --- | --- | --- | --- | --- | --- |
| *cno*Δ*ProxIDR* maternal/zygotic | 6<br>0.19354839 | 10<br>0.32258065 | 8<br>0.25806452 | 7<br>0.22580645 | 31 | 81% |
| *cnoIDRHelixOnly* maternal/zygotic | 18<br>0.2195122 | 23<br>0.2804878 | 33<br>0.40243902 | 8<br>0.09756098 | 82 | 78% |
| *cnoAfadin IDR/cno*<sup>R2</sup> | 18<br>0.51428571 | 7<br>0.2 | 6<br>0.17142857 | 4<br>0.11428571 | 35 | 49% |
| *cnoCanoelDR/cno*<sup>R2</sup> | 31<br>0.88571429 | 3<br>0.08571429 | 1<br>0.02857143 | 0<br>0 | 35 | 11% |
| *cnoIDRΔN/cno*<sup>R2</sup> | 29<br>0.80555556 | 5<br>0.13888889 | 1<br>0.02777778 | 1<br>0.02777778 | 36 | 19% |
| *cnoIDRΔH/cno*<sup>R2</sup> | 52<br>0.88135593 | 7<br>0.11864407 | 0<br>0 | 0<br>0 | 59 | 12% |
| *cnoIDRΔC/cno*<sup>R2</sup> | 21<br>0.47727273 | 13<br>0.29545455 | 5<br>0.11363636 | 5<br>0.11363636 | 44 | 52% |

